# “Tumor Explants Elucidate a Cascade of Paracrine SHH, WNT, and VEGF Signals Driving Pancreatic Cancer Angiosuppression”

**DOI:** 10.1101/2023.03.02.529724

**Authors:** Marie C. Hasselluhn, Amanda R. Decker-Farrell, Lukas Vlahos, Dafydd H. Thomas, Alvaro Curiel-Garcia, H. Carlo Maurer, Urszula N. Wasko, Lorenzo Tomassoni, Stephen A. Sastra, Carmine F. Palermo, Tanner C. Dalton, Alice Ma, Fangda Li, Ezequiel J. Tolosa, Hanina Hibshoosh, Martin E. Fernandez-Zapico, Alexander Muir, Andrea Califano, Kenneth P. Olive

## Abstract

The sparse vascularity of Pancreatic Ductal Adenocarcinoma (PDAC) presents a mystery: what prevents this aggressive malignancy from undergoing neoangiogenesis to counteract hypoxia and better support growth? An incidental finding from prior work on paracrine communication between malignant PDAC cells and fibroblasts revealed that inhibition of the Hedgehog (HH) pathway partially relieved angiosuppression, increasing tumor vascularity through unknown mechanisms. Initial efforts to study this phenotype were hindered by difficulties replicating the complex interactions of multiple cell types *in vitro*. Here we identify a cascade of paracrine signals between multiple cell types that act sequentially to suppress angiogenesis in PDAC. Malignant epithelial cells promote HH signaling in fibroblasts, leading to inhibition of WNT signaling in fibroblasts and epithelial cells, thereby limiting VEGFR2-dependent activation of endothelial hypersprouting. This cascade was elucidated using human and murine PDAC explant models, which effectively retain the complex cellular interactions of native tumor tissues.

## Significance

We present a key mechanism of tumor angiosuppression, a process that sculpts the physiological, cellular, and metabolic environment of PDAC. We further present a computational and experimental framework for the dissection of complex signaling cascades that propagate among multiple cell types in the tissue environment.

## Introduction

PDAC is an aggressive malignancy characterized by a highly desmoplastic microenvironment comprising abundant stromal cells and extracellular matrix (1). This produces a high interstitial fluid pressure that restricts blood flow within the tumor parenchyma, limiting drug delivery while also inducing extreme hypoxia (2–4). Yet, curiously, these conditions do not induce rampant angiogenesis in PDAC as ductal pancreatic tumors are hypovascularized compared to normal pancreatic tissue. Indeed, PDAC exhibits the lowest endothelial index (EI) across 31 cancer types assessed from The Cancer Genome Atlas (TCGA) data (5). The inhibition of angiogenesis under conditions that would typically induce vascular growth, or “angiosuppression”, is an unexplained facet of PDAC biology that nevertheless impacts many aspects of its development, pathophysiology, metabolism, and treatment response.

One potential contributor to PDAC angiosuppression is the HH pathway, which forms a paracrine signal between malignant epithelial cells and nearby cancer-associated fibroblasts (CAFs) (4,6–8). In 70% of PDAC cases (6), the Sonic Hedgehog (SHH) ligand is secreted at high levels from malignant cells, activating downstream signaling in CAFs through binding to the Patched (PTCH1/2) receptors. This relieves inhibition of Smoothened (SMO) leading to the activation of the Glioma-associated Oncogene (GLI) family of transcription factors (9), thus promoting CAF proliferation (7). In prior work, we found that pharmacological inhibition or genetic ablation of SMO in genetically engineered mouse (GEM) models of PDAC led to increased tumor angiogenesis in a VEGFR2-dependent manner (4, 7). However, the mechanism of this effect is unclear as endothelial cells lack active HH pathway signaling and *in vitro* co-culture experiments did not successfully recapitulate the phenotype (7). This experience highlights the challenges of determining molecular mechanisms of complex *in vivo* phenotypes that emerge from the paracrine interactions of multiple communicating cell types.

We approached this challenge in two ways. First, we developed and optimized methods for the short-term culture of intact thick slices of fresh human and murine PDAC. These “tumor explants” maintain the histopathological architecture of the original tumor, with strong representation of the heterogeneous cells present in the PDAC microenvironment. Critically, tumor explants recapitulated the dynamics of angiogenesis instigated by SMO inhibition, serving as a facile system for mechanistic investigations of paracrine cascades.

Second, we leveraged recent developments in the area of regulatory network analysis, a systems biology approach designed to extract mechanistic information from RNA expression data (Supplementary Fig. S1). Regulatory network analysis uses the integrated expression of large sets of genes as multiplexed reporter assays to infer the functional activity of regulatory proteins (e.g. proteins whose function has a large impact on gene expression). This can be performed using very direct regulators, such as transcription factors and chromatin modifiers, where the gene sets are the direct transcriptional targets of the regulatory protein. Alternatively, it can be performed using indirect regulators, such as upstream ligands and receptors, where the gene sets serve as an indirect protein activity signature. In both cases, the gene sets (or “regulon”) for each regulatory protein are generated *experimentally* – in a context-specific manner – using highly validated algorithms based on information theory (10–12). Recent work (13), deployed in the PISCES package (14), has extended this approach for use on single cell RNA sequencing (scRNA-seq) datasets, allowing construction of bespoke regulatory networks for each different cell type present in the tumor. This enables measurements of treatment effects on the activity of most ligands, receptors, and transcription factors in the genome, in each individual cell of a tumor, *in vivo*. The variance stabilization and multiplexing conferred through this approach also largely overcome the limitations of gene dropout that complicate gene expression analysis of scRNA-seq datasets (13).

Using both tumor explants and single cell regulatory network analysis, we found that downstream HH signaling in CAFs initiates a second paracrine signal – secretion of WNT Inhibitory Factor-1 (WIF1) – which can bind the entire family of WNT ligands and prevent their binding to cognate receptors (15–17). Downstream WNT signaling regulates VEGF ligand secretion through established mechanisms (18–21), initiating a third paracrine signal that promotes VEGFR-dependent angiogenesis. Together, these results provide a mechanistic basis for PDAC angiosuppression as a natural consequence of the upregulation of SHH in KRAS-mutant PDAC cells. This also illustrates how cascades of paracrine signals can propagate through tumor tissues to induce complex functional phenotypes, and provides an experimental paradigm for investigating higher order cellular interactions in tissues.

## Results

### WIF1 is a candidate Hedgehog target in PDAC CAFs

Prior studies on the response of murine PDAC to SMO inhibition utilized distinct inhibitors, timepoints, and analytical techniques, drawing divergent conclusions regarding potential effects on angiogenesis (4,7,8). We systematically measured vascularity in PDAC tissues from Kras^LSL.G12D/+^; P53^LSL.R172H/+^; Pdx1-Cre^tg/+^ (KPC) mice treated for varying amounts of time with either the SMO inhibitor IPI-926 (4,7,22) or a vehicle control (hydroxypropyl-beta-cyclodextrin (HPBCD)). Quantification of immunohistochemistry (IHC) for the endothelial cell marker Endomucin (EMCN) revealed that the Mean Vessel Density (MVD) of KPC tumors increased beginning 2 days after IPI-926 treatment, plateauing at 4 days and 7-13 days of SMO inhibition (**Fig. 1A,** Supplementary Fig. 2A). By contrast, and in agreement with a previous report (8), quantification of EMCN-positive pixels per image was not statistically altered upon SMO inhibition (Supplementary Fig. 2B). Next we assessed an early event in angiogenesis, endothelial tip cell formation, using the marker phospho-VEGFR2 (pVEGFR2) (23). Co-immunofluorescence (co-IF) for EMCN and pVEGFR2 revealed a significant increase in endothelial tip cell formation only on day two of SMO inhibition (**Fig. 1B,** Supplementary Fig. 2C,D). These data are best explained by a transient burst of angiogenesis that subsequently elevates the steady-state vascular density following SMO inhibition.

**Figure 1.**
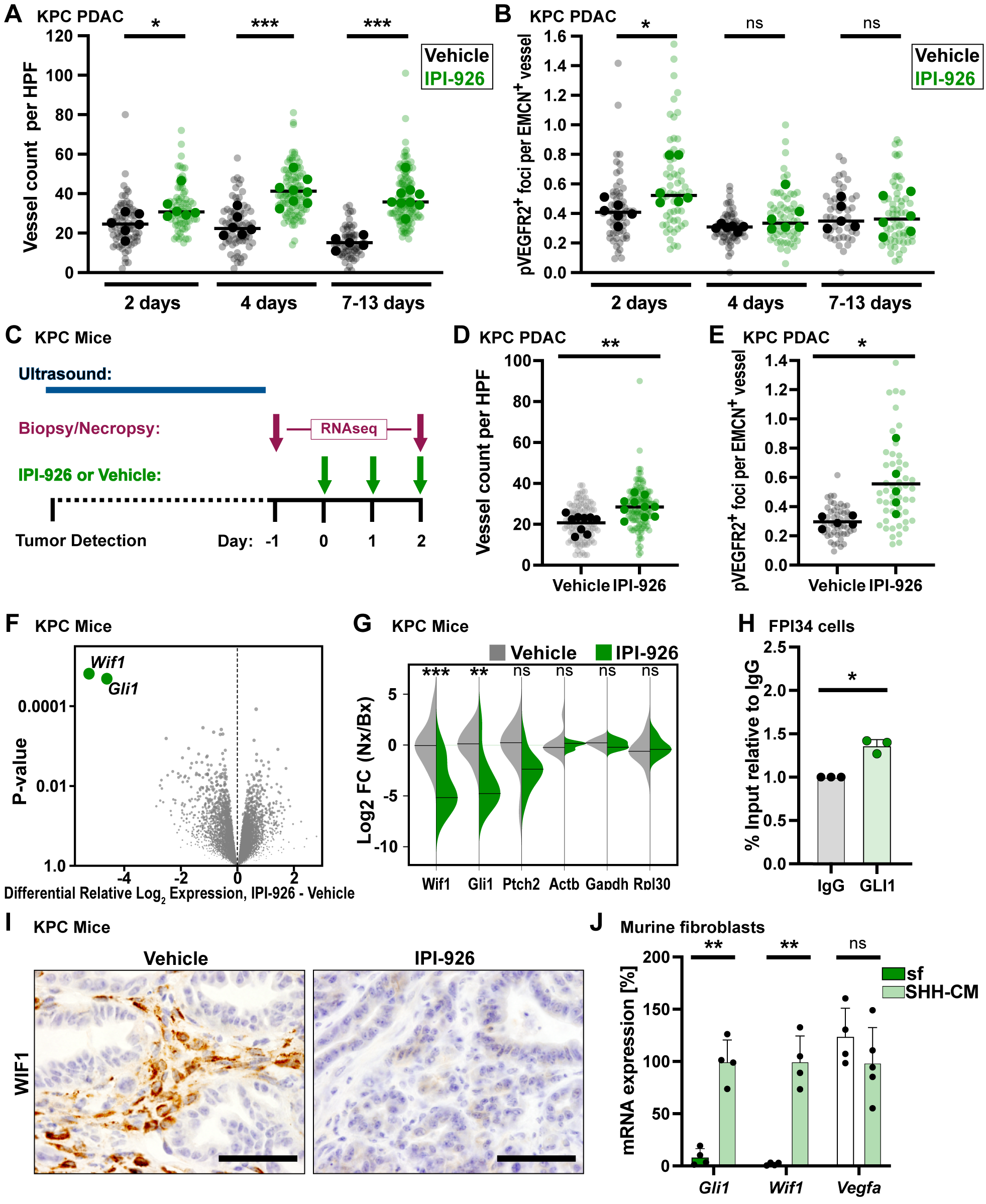
SMO inhibition abrogates SHH-induced WIF1 expression in CAFs. **A,** Tumors from KPC mice treated for the indicated times points with either vehicle or IPI-926 (40 mg/kg) (n=5-8) were stained for the vessel marker EMCN. Quantification of vessel count based on 12 40x fields of view (light dots), averaged per tumor (dark dots), and compared by one-way ANOVA with Tukey correction (*, p<0.05; ***, p<0.001). **B,** Colocalization of pVEGFR2 foci and EMCN as evaluated via co-IF. Quantification of pVEGFR2 foci per EMCN^+^ vessel based on 10 fields of view (light dots), averaged per tumor (dark dots), compared by one-way ANOVA with Tukey correction (*, p<0.05). **C,** Diagram of KPC mouse treatments with vehicle or IPI-926 (40 mg/kg) (n=10 each) for tumor biopsy/necropsy study. **D,** Tumor necropsy samples were stained for EMCN, evaluating mean vessel density (n=9-10). Quantification of vessel count based on 12 fields of view (light shade), averaged per tumor (dark shade), compared by student t-test (**, p<0.01). **E,** Co-IF of pVEGFR2 foci at EMCN^+^ vessels (n=5). Quantification of pVEGFR2 foci per EMCN^+^ vessel based on 10 fields of view (light shade), averaged per tumor (dark shade), compared by student t-test (*, p<0.05). **F,** Significantly regulated genes (green) comparing IPI-926-treated necropsy samples normalized to matching biopsies to HPBCD controls (n=10 each). **G,** Downregulation of HH-responsive genes upon SMO inhibition. Log2 Fold Change of necropsy samples normalized to matching biopsies. Significance indicated (**, p<0.01; ***, p<0.005), based on raw p values, FDR threshold 1.5. **H,** ChIP for GLI1 followed by qRT-PCR on the *WIF1* promoter (n=3) in FPI34 cells, compared by paired t-test (*, p<0.01). Mean and SD are displayed. **I,** Representative image of WIF1 staining in KPC-derived tumors treated with 40 mg/kg IPI-926 for 10 days. Scale = 50 m. **J,** QRT-PCR-based expression analysis of *Gli1*, *Wif1, and Vegfa* in murine fibroblasts in response to treatment with SHH conditioned medium (n=4). Data are normalized to samples treated with SHH-CM. compared by student t-tests (**, p<0.01), mean and SD are shown.

To identify candidate genes or pathways associated with the angiogenic response of PDAC to HH pathway inhibition, we performed an intervention study of IPI-926 in KPC mice harboring pancreatic tumors identified by high resolution ultrasound (24). To control for inter-tumoral heterogeneity, we acquired pre-treatment biopsies via abdominal laparotomy (25) and then randomized mice to treatment with IPI-926 or vehicle (n=10 per group)(**Fig. 1C**). After two days of treatment, mice received a final dose and were euthanized two hours later. MVD and tip cell formation were elevated as expected (**Fig. 1D,E)**. Bulk RNA-seq and differential expression in paired biopsy/necropsy samples in IPI-926-vs. vehicle-treated tumors identified two genes were significantly downregulated: the well-known HH pathway target gene *Gli1* and a WNT pathway inhibitor, *Wif1* (**Fig. 1F,G**, Supplementary Fig. 2E, Supplementary Table 1). WIF1 is a secreted protein that binds to both canonical and non-canonical WNT ligands, preventing their engagement with cognate receptors (26). The *Wif1* promoter harbors canonical GLI binding sites (Supplementary Fig. 2F) and it was previously identified in a signature of genes dysregulated in CAFs sorted from KPC tumors after two weeks of SMO inhibition (8). As the WNT pathway is known to regulate *Vegfa* expression in multiple systems (18–21), we began to investigate its potential role in the response of pancreatic tumors to SMO inhibition.

To validate WIF1 as a candidate GLI target gene in PDAC, we first performed chromatin immunoprecipitation (ChIP) on an immortalized human pancreatic fibroblast line (FPI34) (27) and confirmed direct binding of endogenous GLI1 to the *WIF1* promoter (**Fig. 1H**). Next, we performed IHC for WIF1 on sections of PDAC from KPC mice and observed a stromal pattern of staining that was lost following SMO inhibition (**Fig. 1I**). Indeed, treatment of cultured murine pancreatic fibroblasts with SHH-enriched conditioned medium (SHH-CM) led to significant induction of both *Gli1* and *Wif1* expression. However, *Vegfa* expression in fibroblasts was unaltered in response to SHH-CM (**Fig. 1J**) and further efforts to develop a coculture system that recapitulated the angiogenic response to SMO inhibition were not successful. Together, these data confirm *Wif1* as a direct HH pathway target in PDAC fibroblasts that interferes with the WNT pathway, which is known to modulate angiogenesis. These results also highlight the need for a facile model that can facilitate mechanistic studies of multicellular interactions.

### PDAC explants maintain tissue architecture, viability, and cellular diversity

To better define the mechanism underlying the multicellular interactions in the PDAC microenvironment, we optimized *ex vivo* tumor explants from human PDAC tissue (28–30) and developed a novel protocol for murine PDAC explants (31). Briefly, 300μL fresh slices of either KPC tumors or resected patient samples are cultured on media-soaked gelatin sponge platforms, with a gelatin cover, for up to a week (**Fig. 2A**). Using this approach, tumor slices maintained their histopathological morphology and tissue architecture over time, with ∼75% viability after 5 or 7 days in culture for murine and human explants, respectively (**Fig. 2B,C**). We then performed IHC on formalin-fixed explant tissues to measure markers of proliferation (Ki67) and apoptosis (cleaved caspase 3, CC3). Proliferation rates were stable in human PDAC explants while murine explants demonstrate a modest decrease over time (**Fig. 2D**; Supplementary Fig. 3A,B). For both human and murine explants, the abundance of CC3^+^ cells was unchanged over time (**Fig. 2D;** Supplementary Fig. 3A,B).

**Figure 2.**
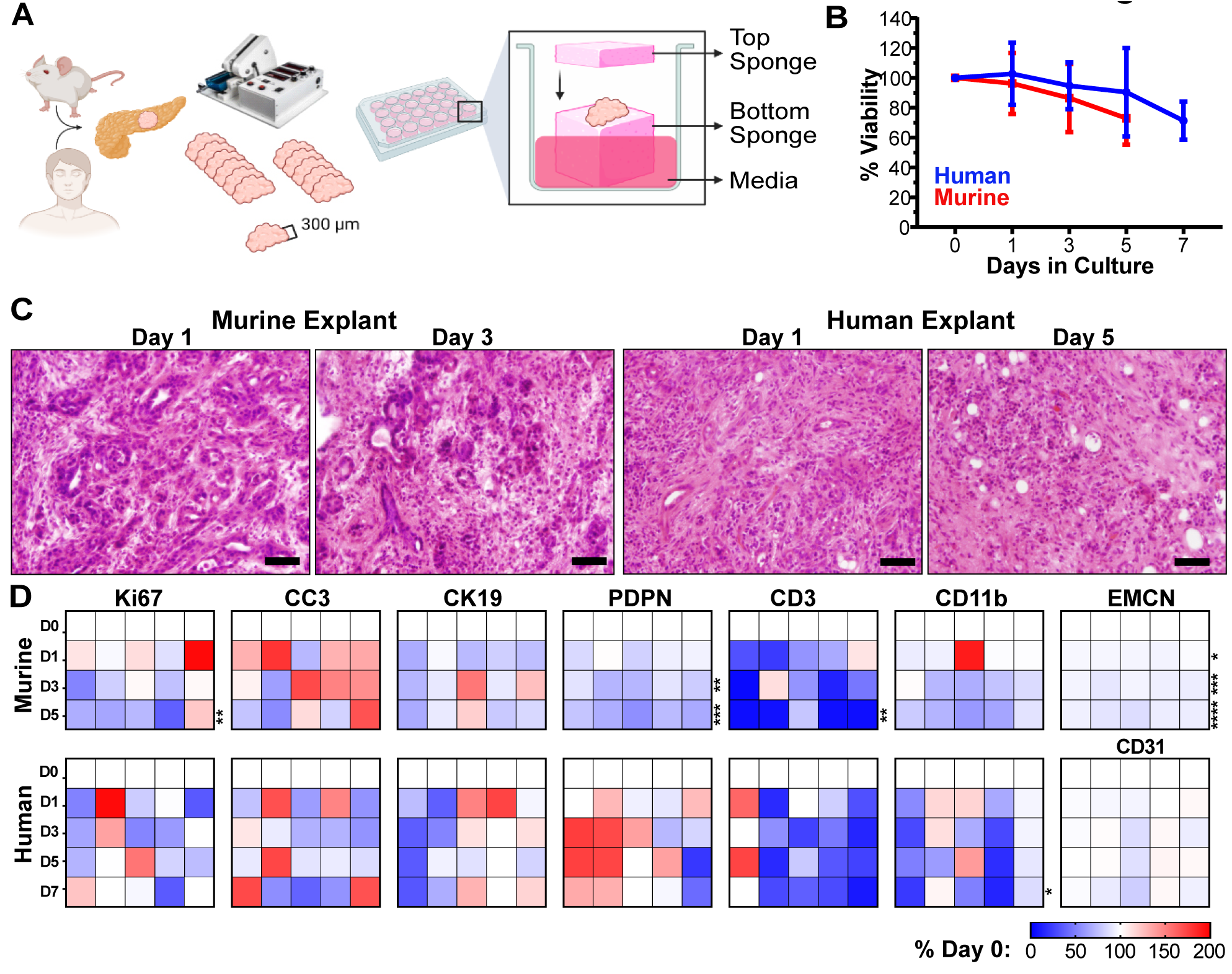
Human and murine PDAC explants maintain tissue architecture, viability, and cellular diversity. **A,** Schematic of human and KPC mouse PDAC explants processing and culturing. **B,** Explant bulk metabolic viability over time as assayed by Alamar Blue (n=5 each). Error bars, SD. **C,** Representative images of Hematoxylin & Eosin (H&E)staining for tissue architecture. Scale = 50 μm. **D,** Murine and human explant time points were stained for various IHC markers for viability (Ki67, proliferation, and CC3, apoptosis) and cell populations (CK19, malignant epithelia; EMCN/CD31, vasculature; Podoplanin, pan-fibroblast; CD3, pan T-cells; CD11b, pan myeloid cells). All quantification time points included day 0, 1, 3 and 5 for murine explants and day 0, 1, 3, 5, and 7 for human explants (n=5 each). Quantification of IHC staining was based on 10-12 fields of view, of which the averaged values per sample per timepoint are represented in the heat maps normalized to day 0 value, compared with two-way ANOVA tests with Dunnett’s correction (*, p<0.05; **, p<0.01; ***, p<0.005; ****, p<0.0001).

Next, to assess whether PDAC explants maintain representation of different cell types throughout the culture process, we quantified individual cellular populations of explants over time, focusing on cancerous epithelia (Cytokeratin 19, CK19), fibroblasts (Podoplanin, PDPN), endothelia (EMCN for murine tissue; CD31 for human tissue), myeloid cells (CD11b), and T cells (CD3) (**Fig. 2D**). We found that the epithelial cell population remained stable in both murine and human explants (Supplementary Fig. 3C). Human CAFs remained stable, while some drop-off was observed in murine explants (Supplementary Fig. 3D). Encouragingly, blood vessel density was remarkably consistent over time, with only a 9% decrease in murine explants at later timepoints (Supplementary Fig. 3E). By contrast, myeloid cells and lymphocytes, which are both normally supplied through peripheral circulation, consistently diminished over time in both murine and human explants (Supplementary Fig. 3F,G). We conclude that explants maintain suitable architecture, viability, and cellular representation, particularly at earlier timepoints in culture.

### SMO inhibition leads to increased angiogenesis via vascular hypersprouting

Although most human and KPC PDAC tumors express high levels of SHH from malignant epithelial cells, most PDAC cell lines express only very low levels of the ligand in 2D cell culture (Supplementary Fig. 4A). We analyzed SHH secretion using a C3H10T1/2 differentiation assay (22) and found that SHH secretion was maintained over the course of 5 days in KPC explants and 7 days in human PDAC explants (Supplementary Fig. 4B,C). Next, in order to assess whether PDAC explants recapitulate the angiogenic response of KPC pancreatic tumors, we treated human and KPC mouse explants with the SMO inhibitors IPI-926 or LDE225, for two or four days (**Fig. 3A-B**). In both models, elevated tip cell formation was observed after two days, followed by an increase in MVD at four days (**Fig. 3A-D**). To ensure the observed angiogenesis was not an off-target effect of high drug concentrations, we performed a dose escalation study with IPI-926 in both KPC and human PDAC explants and found increased endothelial pVEGFR2 beginning at 10nM in both species (Supplementary Fig. 5A,B), consistent with its reported IC50 of 7-10nM (32). Finally, to confirm the specificity of the endothelial pVEGFR2 measurements, we treated KPC explants with the mouse-specific VEGFR2 inhibitor DC101, and human explants with the receptor tyrosine kinase sunitinib, respectively, and observed near-complete loss of endothelial pVEGR2, even in the presence of IPI-926 (Supplementary Fig. 5C,D). These observations validate the ability of PDAC explants to recapitulate dynamic, multicellular phenotypes. They also affirm the effects of SMO inhibition in PDAC using two structurally-distinct agents and demonstrate phenotypic conservation in human PDAC tissue.

**Figure 3.**
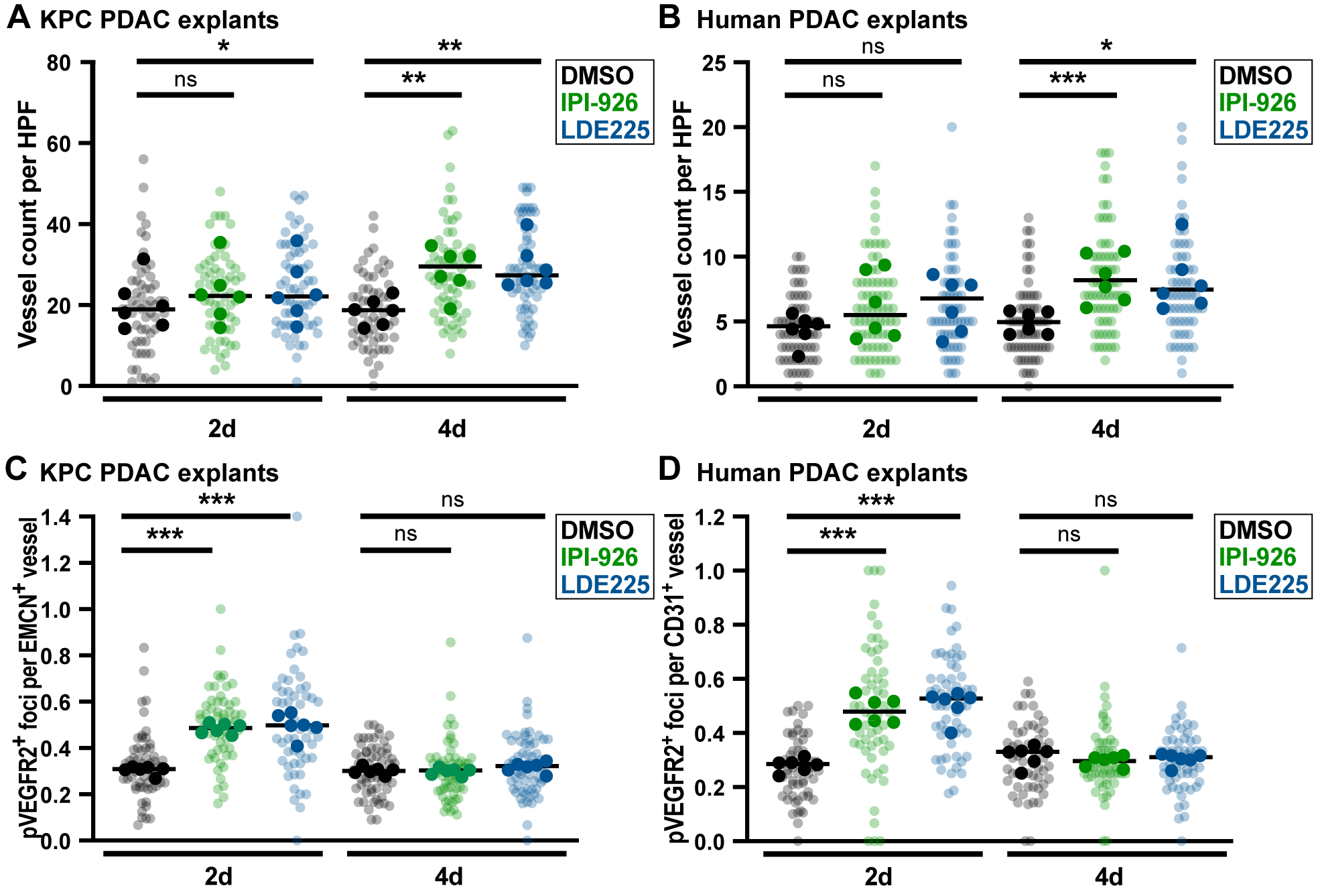
SMO inhibition increases vessel count and induces endothelial hypersprouting in murine and human PDAC explants. **A,** KPC explants treated with DMSO, 1μM IPI-926, or 1μM LDE225, *ex vivo* vessel count using EMCN staining (n=6). Quantification based on 7-12 fields of view (light shade), averaged per tumor (dark shade), compared by one-way ANOVA tests with Tukey correction (*, p<0.05; **, p<0.01). **B,** Human explants treated with DMSO, 1 M IPI-926, or 1μM LDE225, *ex vivo* vessel count using CD31 staining. Quantification based on 7-12 fields of view (light shade), averaged per tumor (dark shade), compared by one-way ANOVA tests with Tukey correction (*, p<0.05; ***, p<0.001). **C,** Co-IF for pVEGFR2/EMCN on KPC explants (n=6). Quantification based on 5-10 fields of view (light shade), averaged per tumor (dark shade), compared by one-way ANOVA tests with Tukey correction (***, p<0.001). **D,** Co-IF for pVEGFR2/CD31 on human PDAC explants (n=6). Quantification based on 5-10 fields of view (light shade), averaged per tumor (dark shade), compared by one-way ANOVA tests with Tukey correction (***, p<0.001).

### WIF1 represses angiogenesis in PDAC

The ability of PDAC explants to model the angiogenic response to SMO inhibition offered a means to study the role of candidate mediators such as WIF1. We therefore treated both KPC and human explants for two days with IPI-926 or LDE225, alone or in combination with recombinant WIF1 protein. In both systems, the restoration of WIF1 through addition of exogenous protein prevented endothelial tip formation, indicating that WIF1 depletion is necessary for the induction of angiogenesis following SMO inhibition (**Fig. 4A,B**). WNT proteins regulate angiogenesis through the induction of *Vegfa* expression via both canonical and non-canonical mechanisms (18–21). Through analysis of public scRNA-seq data (33), we identified *WNT2*, *WNT2B*, *WNT4*, *WNT5A*, *WNT6*, *WNT7A*, *WNT7B*, and *WNT10A* as the most abundant WNT species in human PDAC (Supplementary Fig. 6A). These WNTs are expressed primarily in CAFs, myeloid cells, and malignant epithelial cells, the same cell types that are the primary sources of Vegfa expression in human PDAC (Supplementary Fig. 6B). Indeed, treatment of malignant PDAC epithelial cells, fibroblasts, and macrophages with recombinant WNT5A produced a dose-dependent increase in *Vegfa* expression in both murine malignant epithelial cells and fibroblasts (**Fig. 4C**).

**Figure 4.**
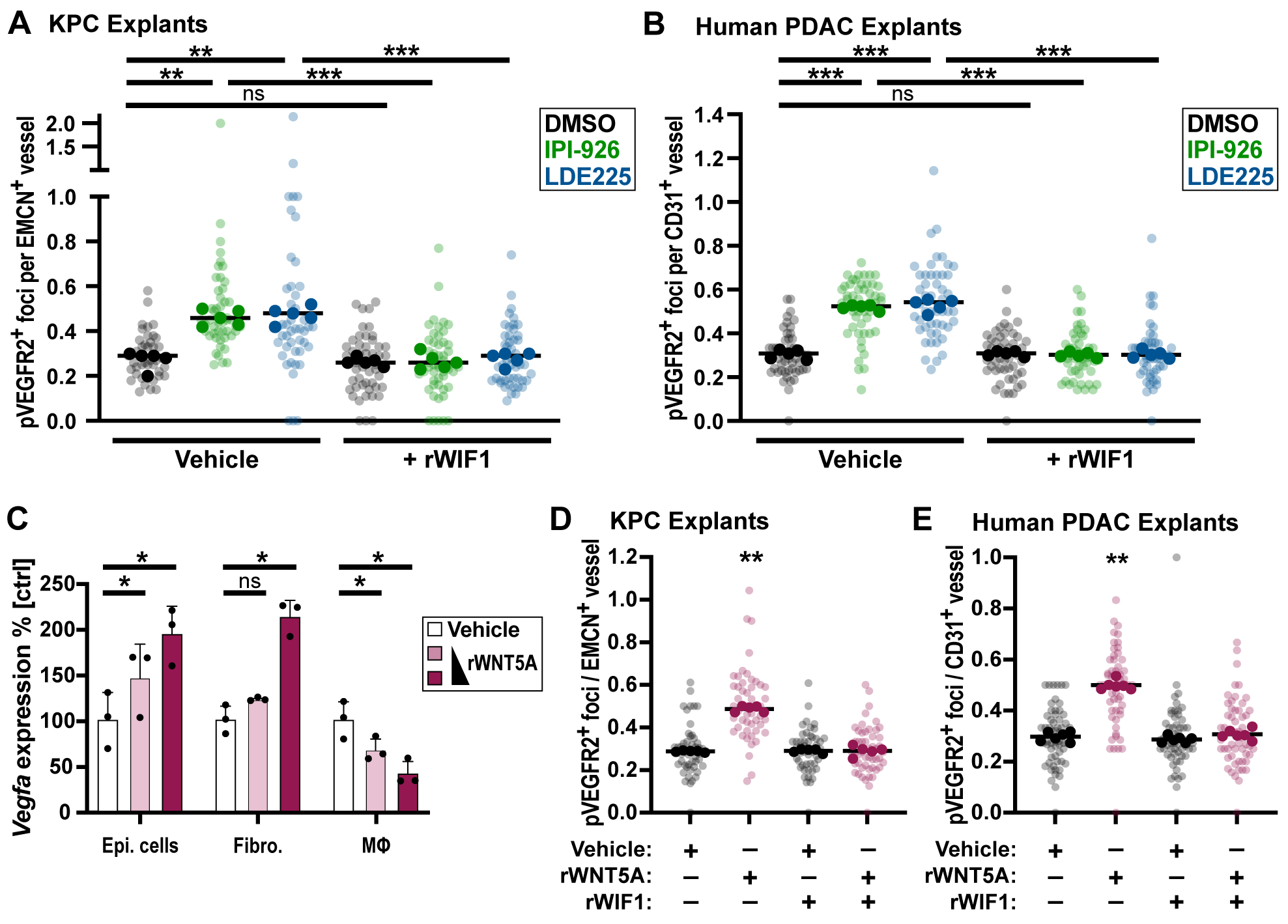
WIF1 blocks WNT5A-induced angiogenesis. **A,** KPC explants treated for 2d with DMSO, 1μM IPI-926, or 1μM LDE225, co-IF for pVEGFR2/EMCN (n=5). Quantification based on 5-10 fields of view (light shade), averaged per tumor (dark shade) compared by one-way ANOVA test with Tukey correction (**, p<0.01; ***, p<0.001). **B,** Human explants treated for 2d with DMSO, 1μM IPI-926, or 1μM LDE225, co-IF for pVEGFR2/CD31 (n=5). Quantification based on 5-10 fields of view (light shade), averaged per tumor (dark shade) compared by one-way ANOVA test with Tukey correction (***, p<0.001).**C,** QRT-PCR for *Vegfa* expression in murine macrophages, epithelial tumor cells, and fibroblasts after 24h treatment with 150ng and 375ng recombinant WNT5A protein (n=3), compared by one-way ANOVA tests with Tukey correction (*, p<0.05). Mean and SD are shown. **D,** KPC explants incubated *ex vivo* under indicated conditions for 2d (750ng rWNT5A, 1 g rWIF1), co-IF for pVEGFR2/EMCN (n=5). Quantification based on 5-10 fields of view (light shade), averaged per tumor (dark shade), compared by one-way ANOVA test with Tukey correction (**, p<0.01). **E,** Human PDAC explants incubated *ex vivo* under indicated conditions for 2d (750ng rWNT5A, 1 g rWIF1), co-IF for pVEGFR2/CD31 (n=6). Quantification based on 5-10 fields of view (light shade), averaged per tumor (dark shade) compared by one-way ANOVA test with Tukey correction for multiple comparisons (**, p<0.01).

To directly test whether WIF1 can regulate angiogenesis via modulation of WNT signaling, we next treated KPC and human PDAC explants with combinations of recombinant WNT5A and WIF1 for two days. Treatment with WNT5A alone increased endothelial pVEGFR2^+^ endothelial tip cell formation in both KPC and human PDAC explants (**Fig. 4D,E**). By contrast, co-treatment with WIF1 reversed the increase in WNT5A-mediated endothelial hypersprouting; administration of WIF1 alone had no effect. We conclude that WIF1 can suppress angiogenesis by inhibiting WNT-dependent activation of VEGFA secretion from malignant epithelial cells and fibroblasts.

### Single cell regulatory network analysis supports a HH-WNT-VEGF cascade regulating PDAC angiosuppression

As an orthogonal means of studying the cascade of paracrine signals in response to SMO inhibition, we performed a treatment experiment in KPC mice and used single cell regulatory network analysis (Supplementary Fig. 7A) to measure the effects of two days of SMO inhibition on the activity of the HH, WNT, and VEGF pathways in PDAC (**Fig. 5A**). Briefly, after pre-processing of the scRNA-seq datasets, we performed Louvain clustering followed by manual refinement to broadly cluster cells type (**Fig. 5B**, Supplementary Fig. 7A). Global shifts in expression were apparent in multiple cell types, indicating a widespread effect from SMO inhibition in PDAC (**Fig. 5C**). As in the earlier biopsy experiment (**Fig.1F-G**), expression of HH pathway target genes such as *Gli1*, *Ptch1*, and *Wif1* were significantly decreased in IPI-926 treated tumors compared to vehicle, an effect that was most apparent in CAFs (**Fig. 5D**).

**Figure 5.**
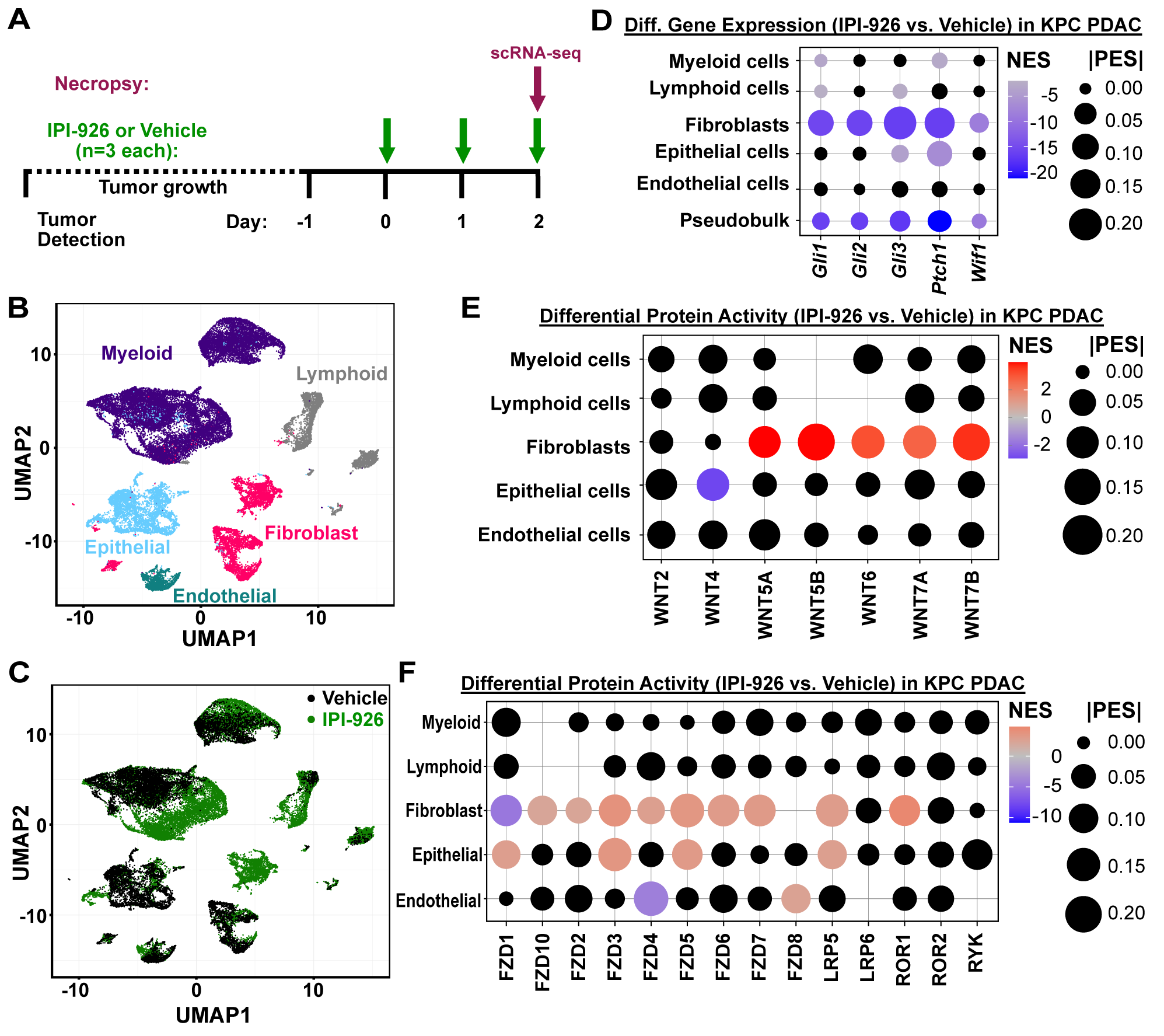
Single cell analyses of KPC pancreatic tumors in response to SMO inhibition. **A,** Diagram of KPC mouse single cell study. Tumor-bearing KPC mice were identified by ultrasound, treated for two days with 40mg/kg IPI-926 or vehicle control, and harvested 2 hours after the final treatment for scRNA-seq of tumor tissues. **B,** UMAP clustering of cells from KPC pancreatic tumors, with cell type assignments. **C**, UMAP clustering of cells from vehicle or IPI-926-treated tumors (n=3 each). **D,** Differential expression of HH-pathway genes comparing IPI-926 to vehicle, in each major cell type. Black dots indicate non-significant differences (p>0.05) according to Mann-Whitney U test. Pseudobulk shows all cells together. **E,** Differential regulatory protein activity analysis shows changes in the inferred activity of WNT ligands, comparing IPI-926 to vehicle, in each major cell type. No dots are displayed for ligands whose activity could not be calculated. Black dots indicate non-significant differences (p>0.05). **F,** Differential regulatory protein activity analysis shows changes in the inferred activity of WNT receptors, comparing IPI-926 to vehicle, in each major cell type. No dots are displayed for receptors whose activity could not be calculated in that cell type. Black dots indicate non-significant differences (p>0.05).

Next we established a computational framework to perform single cell master regulator analysis, using PISCES. Briefly, we first applied ARACNe3 (34) to a collection of PDAC scRNA-seq data from control KPC mice to generate three types of bespoke regulatory networks: one network comprising the inferred direct target genes of ∼1800 transcription factors, cofactors, and chromatin modifiers; a second network comprising indirect functional signatures for ∼2,300 upstream signaling proteins; and a third network comprising indirect functional signatures for ∼1,200 cell surface proteins. This was performed for each major cell type in the tumors, generating sets of context-specific networks for malignant epithelial cells, CAFs, myeloid cells, lymphocytes, and endothelial cells in murine PDAC (Supplementary Table 2). This enabled us to quantify the functional activity of ∼5300 proteins in each individual cell to identify signaling and regulatory changes in response to drug treatment.

We first used this approach to measure changes in the activity of WNT ligands and receptors across cell types in response to IPI-926 treatment. While endogenous WNT levels in human PDAC and vehicle-treated KPC tumors showed expression of a variety of canonical and non-canonical WNTs (Supplementary Fig. 6A, 7B), we consistently observed significant activation of non-canonical WNTs (WNT5A, 5B, 6, 7A, and 7B) in the CAFs of IPI-926 treated, KPC-derived tumors (**Fig. 5E**); changes in WNT ligand activities in other cell types were generally not significant. Similarly, multiple WNT receptors were activated in CAFs as well as in epithelial tumor cells (**Fig. 5F**), consistent with the widespread relief of WNT inhibition due to loss of WIF1 expression.

Finally, we analyzed changes in the expression of angiogenic regulators, including angiopoietin, thrombospondin, and VEGF family members. After 48 hours of treatment with IPI-926, transcription of pro-angiogenic factors has largely been shut down and we observe evidence of up-regulation of anti-angiogenic genes such as thrombospondin-2 (*Thbs2*), particularly in myeloid and epithelial tumor cells (Supplementary Fig. 7C). THBS2 counteracts VEGF-induced angiogenesis and often serves as a feedback response that limits bursts of angiogenesis (35, 36). These observations are consistent with the rapid loss of pVEGFR2 expression by four days of Smo inhibition, in both explants and KPC mouse pancreatic tumors (**Fig.1B, Fig. 3C,D**), and help explain why angiogenesis stabilizes at an elevated threshold rather than continuing unchecked.

Taken together, our findings using human and murine PDAC explants, human datasets, and GEM models detail a cascade of three paracrine signals that propagate between multiple cell types and collectively serve to limit angiogenesis in PDAC (Supplementary Figure 7D). Oncogenic KRAS activation leads to increased expression and secretion of SHH from malignant epithelial cells, leading to paracrine activation of GLI transcription factors in CAFs. GLI genes induce WIF1 expression and secretion, thereby restraining the activation of VEGF signaling by downstream WNT signaling in multiple cell types. Conversely, SMO inhibition releases the pro-angiogenic activity of WNTs, particularly through activation of non-canonical WNT receptors, leading to a burst of VEGFR2 activation in endothelial tip cells, an effect that is quickly counteracted through upregulation of THBS2.

## Discussion

The expansive desmoplastic stroma of PDAC is a pathognomonic feature of this complex and deadly disease. Though driven indirectly by mutations in malignant epithelial cells, once established the tumor microenvironment broadcasts a cacophony of intercellular signals, with putative communication between every possible pair of cell types (37). Two decades of laborious effort have helped elucidate numerous individual paracrine signaling pathways that mediate communication between individual pairs of cell types in PDAC. Our findings clarify that these signals do not stop at the target cell. Instead, they propagate a cascade of signals that ripple out from every cell, interacting, interfering, and ultimately sculpting an ecosystem that is robust to disruption – a natural homeostasis that likely contributes to the extraordinary therapeutic resistance of PDAC.

Here we provide an investigative and analytical framework for studying the higher order complexity of paracrine cascades. Co-culture models using isolated cell types or organoids have proven invaluable for the study of individual paracrine signals between pairs of cells. However, the dissociation of tumor tissues destroys the native complexity and spatial structure of the tissue. Elements may be reconstituted, but it is not currently possible to fully restore PDAC tissue from constituent parts. Instead, we set out to preserve the complexity of PDAC tissues, building on the work of prior efforts with human PDAC (28–30) and extending them to include murine PDAC. The resulting models and media, which incorporate information on the metabolic composition of PDAC interstitial fluid (38), are suitable for short-term experiments with small molecule drugs, blocking antibodies, recombinant proteins, and other perturbations to modulate cell biology over the course of hours or days. Sandwiching the explants in media-infused gelatin also protects the tissue from high atmospheric oxygen levels and creates an artificial gradient of nutrients and waste that may mimic aspects of PDAC physiology. The availability of both murine and human PDAC model systems enables direct comparisons of mechanisms and drug effects across species – a key component of preclinical translation. While these and prior version of PDAC explants are limited due to attrition of cells derived from peripheral circulation (30), we anticipate future iterations that are supplemented with matched immunocytes or incorporated into bioengineered “organs on a chip” (39) to further refine the system.

PDAC explants were instrumental in our efforts to explore the mechanisms of angiosuppression – the confounding deficit neoangiogenesis in PDAC under highly hypoxic conditions. While there are undoubtedly additional contributors to this phenotype, our data highlight a cascade of three paracrine pathways – HH to WNT to VEGF – as a major suppressor of angiogenesis. The activation of HH signaling through upregulation of SHH ligand expression in malignant epithelial cells appears to be a consequence of KRAS mutation, though the mechanism is unknown. By tracing the path from HH to VEGFR2, this aspect of angiosuppression is established a natural consequence of KRAS mutation, reflecting the fact that tumor evolution is anchored to preexisting genetics and regulatory environment. The mechanism also implicates WNT signaling as a key regulator of angiogenesis in PDAC, adding to its recently-discovered role in immunosuppression (37). We show that WNT-mediated modulation of angiogenesis is titrated by the HH pathway via regulation of WIF1, a potent suppressor of WNT ligand function (40). Given the highly pleiotropic family of WNT ligands expressed from multiple cell types in PDAC, WIF1 serves a key choke point on the activity of the entire pathway.

The complexity of WNT and other pathways, which include dozens of ligands, receptors, transducers, and transcription factors, highlights the importance of computational techniques as a complement to experimental manipulation of individual factors. While single cell gene expression analysis has begun to enable the description of cell types in complex tissues, analytical challenges such as gene dropout from low read depth complicate efforts to trace molecular biology mechanisms at the individual cell level. Single cell regulatory network analysis largely overcomes many of these limitations, enabling the experimental measurement of pathway activity in individual cells of intact tumors in response to drug treatments or other perturbations. Thus, in addition to measuring the effects of adding a single WNT ligand (WNT5A) to explants, we could also measure the changes in activity of the entire family of WNT ligands and receptors following SMO inhibition in GEM. These complementary approaches establish an investigative framework for understanding complex phenotypes in intact tissues.

## Summary

Pancreatic tumor explants reproduced the complex phenotype of angiosuppression in PDAC and facilitated mechanistic dissection of contributing pathways. Combined with single cell regulatory network analysis, we elucidated a cascade of three paracrine pathways bridging between multiple cell types, that connect KRAS mutation to angiosuppression via HH, WNT, and VEGF signaling.

## Material and methods

### Animal Breeding, Enrollment, and Dosing

All animal research experiments were approved by the Columbia University Irving Medical Center (CUIMC) Institutional Animal Care and Use Committee. Mouse colonies were bred and maintained with standard mouse chow and water, *ad libitum*, under a standard 12hr light/12hr dark cycle. KPC (*Kras*^LSL.G12D/+^; *P53*^LSL.R172H/+^; *Pdx1-Cre*), KC (*Kras*^LSL.G12D/+^; *Pdx1-Cre*), PC (*P53*^LSL.R172H/+^; *Pdx1-Cre*) mice were generated in the Olive Laboratory by crossing the described alleles. Mouse genotypes were determined using real time PCR with specific probes designed for each gene (Transnetyx; Cordova, TN).

KPC mice were monitored by manual palpation for tumor development, confirmed via ultrasound, and included in studies when tumors reached dimensions between 4-6 mm. Enrolled mice were then randomized to study arms. *Post hoc* analysis determined no significant enrichment for sex in any arm of the studies was observed. Treatment with the vehicle hydroxypropyl-beta-cyclodextrin (HPBCD, Acros Organics; 5% w/w in water for injection (WFI)), or IPI-926 (kindly provided by PellePharm; 5 mg/ml) was performed daily via oral gavage at 40 mg/kg for the indicated time points (2 days, 4 days, or 7-13 days).

### Histological Stainings: Immunohistochemistry (IHC) and Immunofluorescence (IF)

4 μm formalin-fixed paraffin-embedded sections were rehydrated using a Leica XL ST5010 autostainer. Slides were subjected to heat-activated epitope retrieval and IHC slides underwent quenching of endogenous peroxidases prior to incubation with primary antibodies (supplementary table 3). For IHC, secondary antibody incubation and development with DAB was followed by hematoxylin counterstain before dehydration and coverslip mounting. IF slides were incubated with fluorochrome-coupled secondary antibodies prior to DAPI staining (Biolegend, 422801) and mounting. Quantitative analyses of IF and IHC images were performed using Fiji (41).

### Differential Gene Expression of KPC Bulk RNA-seq Data

Short-term intervention studies using IPI-926 or vehicle (HPBCD) were performed in tumor-bearing KPC mice. We acquired pre-treatment biopsies as previously described (25) before randomizing mice into respective treatments for two days. Matching biopsy and necropsy samples were subjected to bulk RNA-seq. To contrast both within subjects, i.e. necropsy vs. biopsy samples, and between treatments, we leveraged a generalized linear model (GLM) as implemented in the edgeR R package (42) using raw count data. First, we adjusted for baseline differences between the mice by initializing the design matrix considering mouse identifiers. Next, we defined treatment-specific necropsy effects and appended them to the design matrix. After estimating the dispersions, we fit the GLM and contrasted the treatment-specific necropsy effects to find genes that behave differently between necropsy and biopsy in vehicle-treated vs. IPI-926-treated mice.

### Chromatin Immunoprecipitation (ChIP)

ChIPs were conducted as previously described (43). Briefly, FPI34 cells (10×10^6^) were cross-linked with 1% formaldehyde, followed by cell lysis. DNA was sheared with sonication for 35 cycles (30-s on/off cycles) in a Diagenode Biorupter 300, and aliquots of the sheared chromatin were then immunoprecipitated using magnetic beads and corresponding antibodies (GLI1: NB600-600, Novus Biologicals, RRID:AB_2111758; IgG: ab18443, Abcam, RRID:AB_2736846). Following immunoprecipitation, cross-links were removed, and immunoprecipitated DNA was purified using spin columns and subsequently amplified by quantitative PCR. PCR primers were designed to amplify a region of the WIF1 promoter containing potential GLI1 binding sites. QRT-PCR was performed in triplicate for each sample using the C1000 Thermal Cycler. Results were represented as % input relative to IgG, where each antibody signal was normalized to its respective input and then relative to the nonimmune IgG control signal.

### RNA isolation and qRT-PCR

Primary tumor cells derived from the KPC GEMM, fibroblasts and myeloid cells were cultivated in DMEM supplemented with 10% fetal bovine serum and 1x penicillin/streptomycin. Cells were seeded in 6-well plates, treated with indicated concentrations of WNT5A (R&D systems, 645-WN) the next day and RNA was harvested after 24h treatment using TRIzol (Thermo Fisher Scientific, 10296010). Subsequent to RNA isolation, cDNA was transcribed using iScript cDNA Synthesis Kit (Bio-Rad, 1708891) and qRT-PCR was performed using Itaq Universal SYBR (Bio-Rad, 1725122) on a StepOne Real-Time PCR system (Applied Biosystems) using listed primer sequences (supplementary table 4). Data were analyzed using normalization to the house keeping gene *Rplp0* via the ΔΔCT approach.

### Explant Sponge Preparation

Powdered porcine gelatin (Sigma Aldrich, G2500) and deionized water were combined to form a 6% w/v solution and gently mixed at 60°C until fully dissolved. The solution was then whisked with a hand-mixer at room temperature until well aerated and stiffened into peaks. On a clean metal tray, a 1cm x 1cm x 1cm silicone mold (Amazon, B07PWPCD34) was pushed into the gelatin mixture until flush with the tray surface. The gelatin and mold are lyophilized in a freezer dryer (supplementary table 5). Dried bulk sponge and mold was transferred onto a silicone mat and baked in a convection oven at 300°F (160°C) for 3 hours to cross-link polymers. Completed sponges were removed from the mold and trimmed to a uniform 1cm cube with a sterile scalpel, then stored in an air tight glass jar with a desiccant packet at room temperature.

### Explant Media Composition and Preparation

Explant media was prepared in a sterile environment, either in a tissue culture hood or on the benchtop with a Bunsen burner flame. Concentrated stock solutions for all components were prepared and stored according to manufacturer’s instructions. In a clean and sterile autoclaved flask, species-specific components (28, 44), select organoid essentials (45), metabolic supplements, pancreas supplements (30), and anti-TIF supplements (38) were combined (supplementary table 6). Media was then filtered into 50 mL aliquots using a vacuum filtration system (0.22 μm filter) and stored at 4°C for up to a month.

### Explant Tissue Collection and Sectioning

Murine tumors were collected following humane euthanasia and trimmed of healthy pancreas tissue in a sterile petri dish. Human tissue samples were obtained from de-identified patients undergoing resection surgeries, primarily pancreaticoduodenectomy (Whipple) or distal pancreatectomy, at New York-Presbyterian/Columbia University Irving Medical Center. Bulk resected tissue was processed by Department of Pathology and tumor tissue was placed into cold DMEM and transported on ice. All tumor tissue was embedded in 2.5% agarose and sectioned into 300 μm slices using a Compresstome. Tumor slices were immediately transferred into ice-cold Hank’s Balanced Salt Solution and kept on ice until plating. Any tumor tissue remaining after sectioning was fixed in 4% paraformaldehyde (Santa Cruz Biotechnology, sc-281692) for 2 hours, at 4°C as the Day 0 control. Fixed tissue was then transferred to 70% ethanol and paraffin embedded for long-term storage.

### Explant Metabolic Viability Assay

Individual explants were weighed prior to plating on Day 0. At each time point, explants were transferred directly into a new 24-wells with 500 μL fresh DMEM (Gibco Life Technologies, 12430062) or RPMI 1640 (Gibco Life Technologies, 21870-076) media (for mouse or human tissue respectively) and 50 μL Alamar Blue (BioRad, BUF012B), with a corresponding media only control well, and incubated at 37°C, 5% CO_2_ for 4 hours. Following incubation, 100 μl of each sample was transferred to a 96-well plate (Corning, 3603) in triplicate and fluorescence (ex. 560 nm, em. 590 nm) was measured on a Varioskan LUX Microplate Reader. For analysis, background levels were subtracted from raw results, and were then normalized to first represent fluorescence per initial tissue weight, then further normalized to be represented as a percentage of Day 0 signal/weight. Five independent samples were evaluated, with at least two explants per time point and three technical replicates per sample.

### Explant Culture and *ex vivo* Treatment Conditions

Gelatin sponges (1cm^3^) were incubated in 24-well plates with 750 μL of respective media at 37°C for at least 30 minutes to soak. According to the respective treatment condition, media was supplemented with DMSO (ctrl; Fisher Bioreagents, BP231-100), IPI-926 (SMO inhibitor; PellePharm), LDE225 (SMO inhibitor; ChemieTek, CT-LDE225), IgG (InVivoMab rat IgG1 isotype control, anti-horseradish peroxidase; Bio X Cell, BE0088; RRID: AB_1107775), α VEGFR2 (InVivoMab anti-mouse VEGFR2 (DC101); Bio X Cell, BE0060; RRID: AB_1107766), Sunitinib (Selleck Chemicals, S7781), recombinant human WIF1 (R&D systems, 1341-WF-050/CF), recombinant murine WIF1 (R&D systems, 135-WF) or recombinant WNT5A (R&D systems, 645-WN). Sectioned explants were transferred to sponges and flattened with forceps and metal spatula, covered with a thin (2-3 mm thick) gelatin top sponge, and incubated in standard cell culture conditions (37°C, 5% CO_2_). Media was replaced daily with 500 μL fresh media. Explants were fixed in 4% paraformaldehyde for 2 hours at 4°C, then transferred to 70% ethanol and paraffin embedded for long-term storage.

### Single Cell Preparation and Sequencing

For single cell RNA sequencing (RNA-seq), respective KPC mice were treated with HPBCD or IPI-926 for 2 days. 2 hours following the final dose, tumor tissue was collected following humane euthanasia and trimmed of healthy pancreas tissue in a sterile petri dish. The tumor pieces were dissociated using a modified protocol based on Miltenyi (mouse) Tumor Dissociation Kit (Miltenyi Biotec, 130-096-730). Briefly, the tumor tissue was placed in a digestion buffer containing trypsin, DNase, and an enzymatic cocktail (supplementary table 7) and digested at 37°C for 42 minutes (37C_m_TDK_2 program) on a gentleMACS Octo Dissociator (Miltenyi Biotec, 130-096-427). Cell suspensions were then filtered through a 40 μm cell strainer (Corning, 431750) and red blood cells were removed by incubation with red cell lysis buffer (Millipore Sigma, 11814389001), before a final resuspension in 100 μL PBS + 0.01% BSA. Samples were submitted to the Sulzberger Genome Center for analysis. Briefly, single-cell sequencing data were processed using the Cell Ranger pipeline (v.3) from 10X GENOMIC. FASTQ files were aligned on gex-mm10-2020-A transcriptomes. All the count matrices were filtered for low quality cells, normalized to CPM and analyzed independently.

### ScRNA-seq Quality Control and Batch Combination

ScRNA-seq profiles from each of the six samples were quality controlled based on minimum and maximum reads per cell in UMIs (Min / Max Depth), the maximum number of unique genes detected (Max. Genes), and the percentage of mitochondrial reads (Max MT%). Parameters were fit to each sequencing dataset individually, with thresholds given (supplementary table 8). Data were inspected for extreme batch effects using a principal component analysis (PCA) in gene expression space. Since no dramatic differences were observed, we concluded the data could be combined with appropriate integration. Single-cell profiles from the six samples were combined using the Seurat scRNA-seq integration protocol (46). In order to preserve more features for subsequent protein activity inference, we adjusted the number of integration features to 4000 (nfeatures).

### Cell Type Mapping

The integrated scRNA-seq dataset was clustered in gene expression space using the standard Seurat SCTransform procedure outlined in (47). Clusters were then mapped to cell types based on the expression of selected markers as well as inspection of unsupervised cluster markers as identified by Seurat’s ‘FindMarkers’ function. Cell type identifiers were manually curated. Because our questions focused on stromal compartments, we did not extensively investigate the difference between malignant and normal epithelial cells in these samples. Notably, InferCNV (48) analysis was inconclusive in terms of identifying clearly mutated populations of epithelial cells, leading us to analyze the entire epithelial compartment as one unit.

### Differential Expression Analysis

Within each cell type, data were re-integrated across samples using the same procedure described previously. For each gene in the resulting integrated, normalized matrix, a Mann-Whitney U-Test (49) between cells in the vehicle and IPI-926 conditions. Rank biserial correlation (RBSC) was reported as the effect size, while p-values were corrected with the Benjamini-Hochberg method (50).

### Protein Activity Analysis

Protein activity was inferred using the PISCES pipeline (14). Regulatory networks were inferred in a cell-type specific manner using ARACNe3 (34). Cells from across samples were pooled within each cell type and subset to 1,000 samples for network generation. The following ARACNe3 parameters were used; 100 subnetworks, 0.25 FDR, no metacells. Protein activity was inferred for each cell type using the appropriately matched regulatory network. Gene expression signatures were generated in the manner described previously (see Differential Expression Analysis), with the p-value transformed to a normalized enrichment score (NES) using ‘pnorm’ and the sign determined by the sign of the RBSC. For each cell type, this created a single signature vector – one value for each gene – within each cell type for the comparison between IPI-926-treated cells and vehicle-treated cells. Activity was then inferred using NaRnEA (34). This produces a NES – a measure of the statistical significance – and a proportional enrichment score (PES) – a measure of effect size. NES values were transformed to p-values using the ‘qnorm’ function, then corrected for multiple hypotheses using the Benjamini Hochberg procedure.

### Visualizations

Heatmaps were generated using the ComplexHeatmap package in R (51). All other plots (scatter plots, dot plots, bar graphs) were generated using ggplo2 in R (52).

### Data Availability Statement

The data generated in this study are available upon request from the corresponding author.

### Ethics Statement

All animal research experiments were approved by the Columbia University Irving Medical Center (CUIMC) Institutional Animal Care and Use Committee (IACUC). All work using patient samples were performed with approval from the CUIMC Institutional Review Board (IRB).

## Supporting information

All Supplemental Tables

## Acknowledgements

This study was funded by NIH grants 1U01CA274312 and 2R01CA215607 (KPO), a Lustgarten Foundation Clinical Translational Program Award (KPO). MCH received support from Deutsche Forschungsgemeinschaft (439440500). ARDF received support from NIH F31-5F31CA250443. ACG received support from the Charles H. Revson Senior Fellowship in Biomedical Science (Grant No. 22-22). MEFZ was supported by CA265050 and Mayo Clinic Cancer Center. This work was supported by the HICCC Cancer Center Support Grant (P30CA013696) including use of the following shared resources: Confocal and Specialized Microscopy, Genomics and High Throughput Screening, Oncology Precision Therapeutics and Imaging, and Molecular Pathology. We thank Marina Pasca di Magliano and Ben Allen for helpful discussions as well as Nina Steele for technical advice and provision of the fibroblast cell line. We are grateful to members of the Olive laboratory for their comments and suggestions.

## Supplementary Figure 1

**Supplementary Figure 1.**
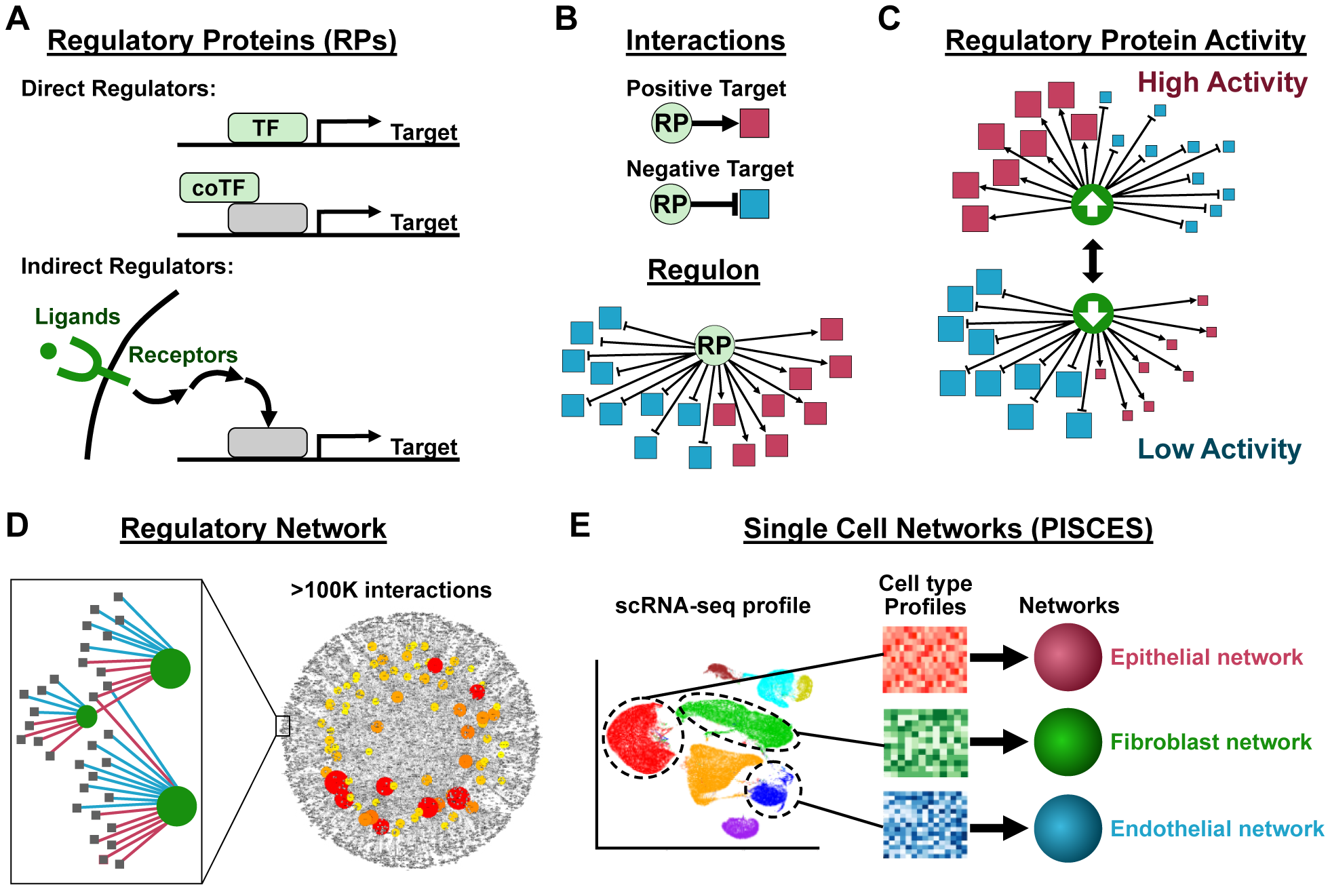
Overview of single cell regulatory network analysis. (**A**) Regulatory proteins include both direct regulators such as transcription factors and co-factors, as well as indirect regulators such as upstream ligands and receptors. (**B**) Each regulator modulates sets of target genes, including positive (transactivating) targets and negative (transrepressing) targets. The total set of target genes for a regulatory protein is called a regulon. (**C**) The activity of a regulatory protein is calculated from the relative expression of its positive and negative targets. (**D**) The sets of target genes for each regulatory protein in the genome can be predicted, with high accuracy, from large numbers (>100) of gene expression profiles for a given biological entity. The predicted regulons are *context specific*, meaning different target genes will be predicted for the same regulatory protein in different tissues or cell types. These predictions are made using the highly validated ARACNe algorithm. (**E**) The PISCES framework is optimized for generating networks using single cell data, such that different context-specific networks are constructed for major cell type in the tumor.

**Supplementary Figure 2.**
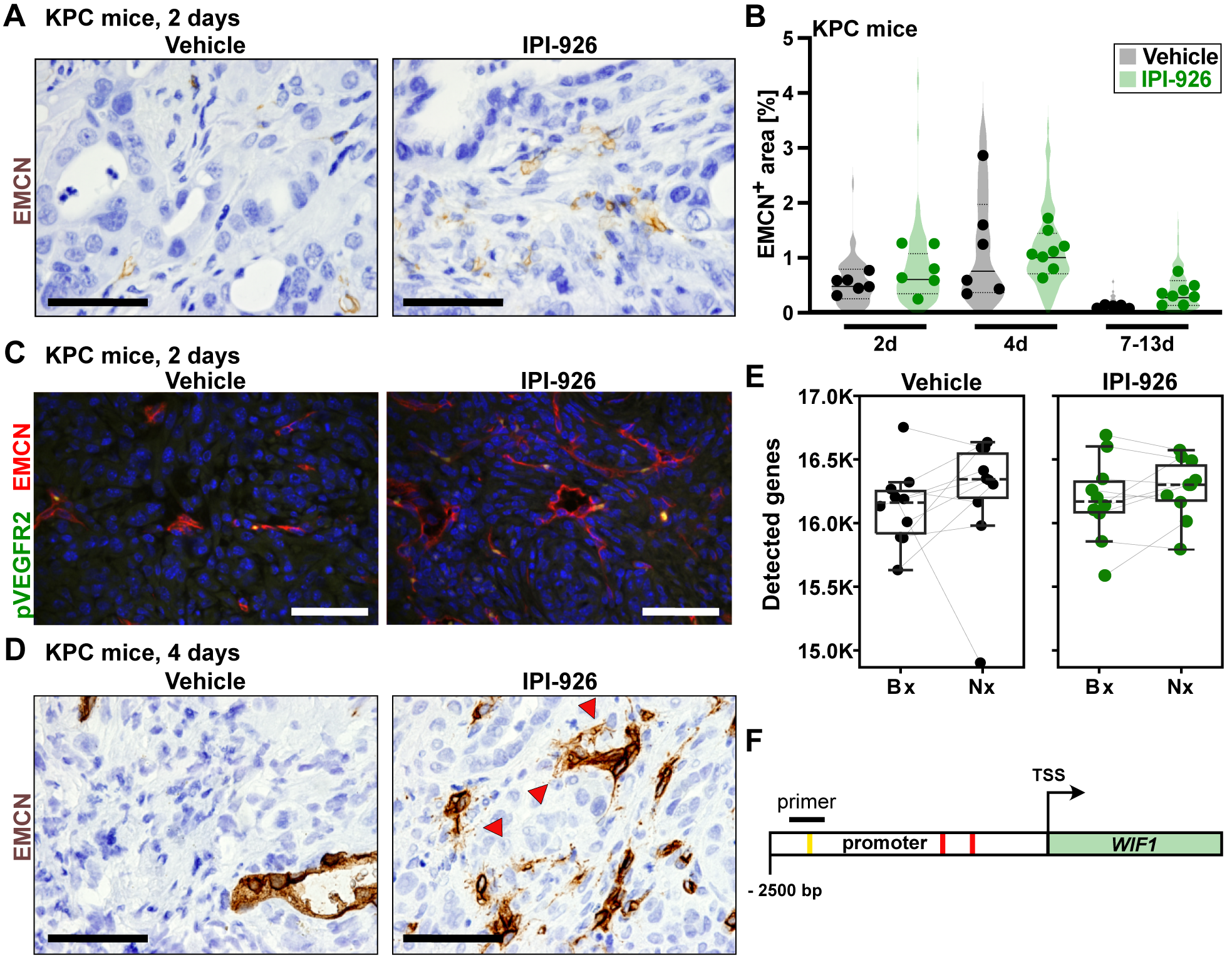
IPI-926 treatment alters cellular composition in the tumor microenvironment. (**A**) Representative images for EMCN staining in KPC mice after 2d of vehicle or IPI-926 treatment. Scale bars, 50 μm. Objective, 100x. (**B**) KPC *in vivo* EMCN area analysis at different time points (n=5-8). Quantification based on 12 fields of view (light shade), averaged per tumor (dark shade). (**C**) Representative images for EMCN/pVEGFR2 co-IF in KPC mice after 2d of vehicle or IPI-926 treatment. Scale bar, 50 μm. Objective, 40x. (**D**) Representative images for EMCN staining after 4d of vehicle or IPI-926 treatment. Red arrowheads indicate tip cell filopodia. Scale bar, 50 μm. Objective, 100x.(**E**) Number of detected genes in RNA-seq libraries from experiment in Fig. 1C. (**F**) Schematic of the WIF1 promoter site, indicating GLI1 binding sites and ChIP primer location.

**Supplementary Figure 3.**
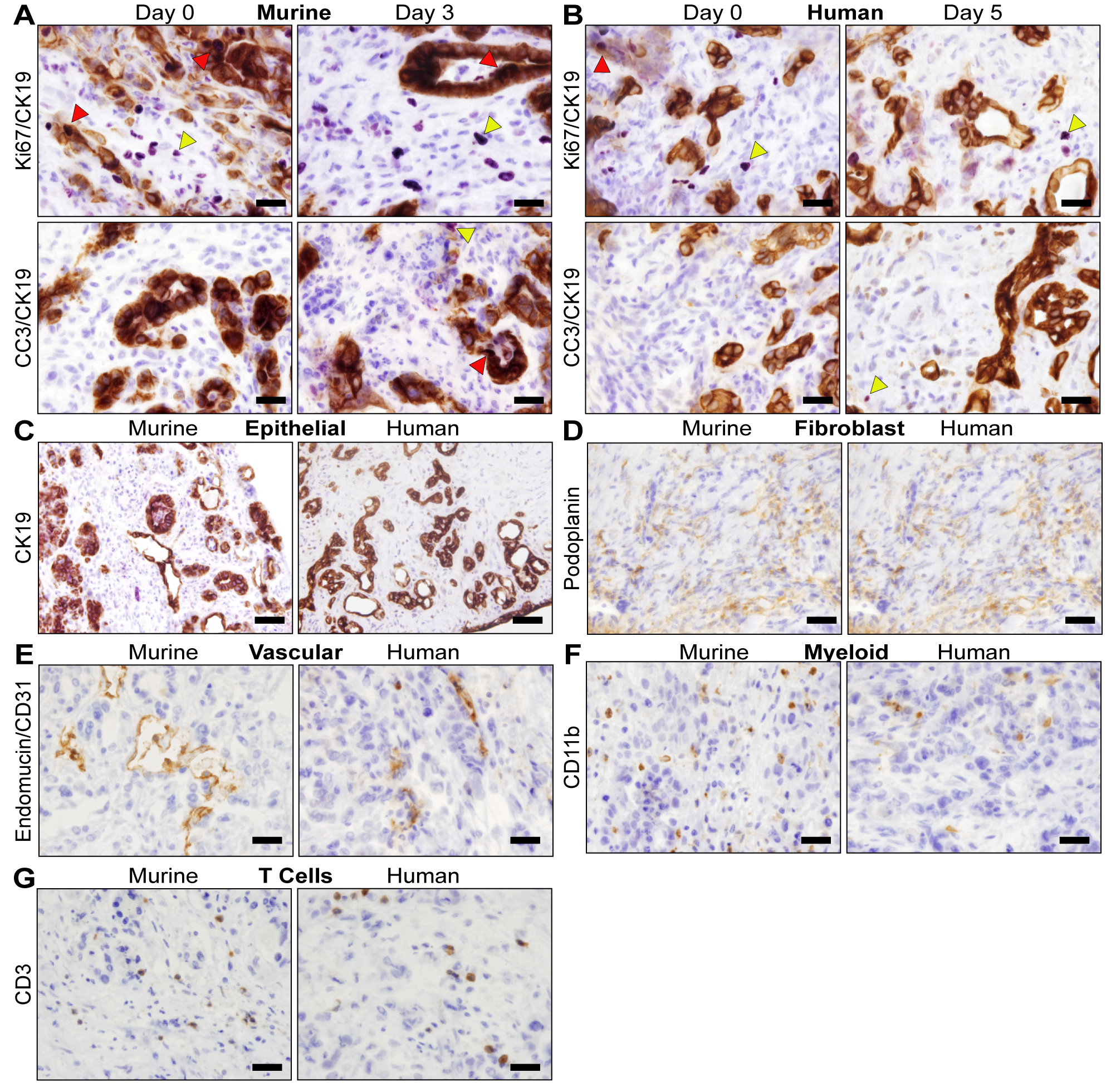
Explant Cellular Populations. (**A**) Representative images for dual IHC staining in murine PDAC explants for malignant epithelial cells (CK19, brown) and proliferating cells (Ki67, purple) or apoptotic cells (CC3, purple) at two timepoints during the culturing process. Red arrows indicate Ki67 or CC3 positive epithelial cells, yellow arrows indicate positive stromal cells. Scale bars, 20 μm. Objective, 100x. (**B**) Representative images for dual IHC staining in human PDAC explants for malignant epithelial cells (CK19, brown) and proliferating cells (Ki67, purple) or apoptotic cells (CC3, purple) at two timepoints during the culturing process. Red arrows indicate Ki67 or CC3 positive epithelial cells, yellow arrows indicate positive stromal cells. Scale bars, 20 μm. Objective, 100x. (**C**) Representative images for CK19 staining in murine and human PDAC explants, for malignant epithelia. Scale bars, 50 μm. Objective, 40x. (**D**) Representative images for podoplanin staining in murine and human PDAC explants, for fibroblasts. Scale bars, 20 μm. Objective, 100x. (**E**) Representative images for ECMN or CD31 staining in murine and human PDAC explants, respectively, for endothelial vasculature. Scale bars, 20 μm. Objective, 100x. (**F**) Representative images for CD11b staining in murine and human PDAC explants for myeloid cells. Scale bars, 20 μm. Objective, 100x. (**G**) Representative images for CD3 staining in murine and human PDAC explants for T-cells. Scale bars, 20 μm. Objective, 100x.

**Supplementary Figure 4.**
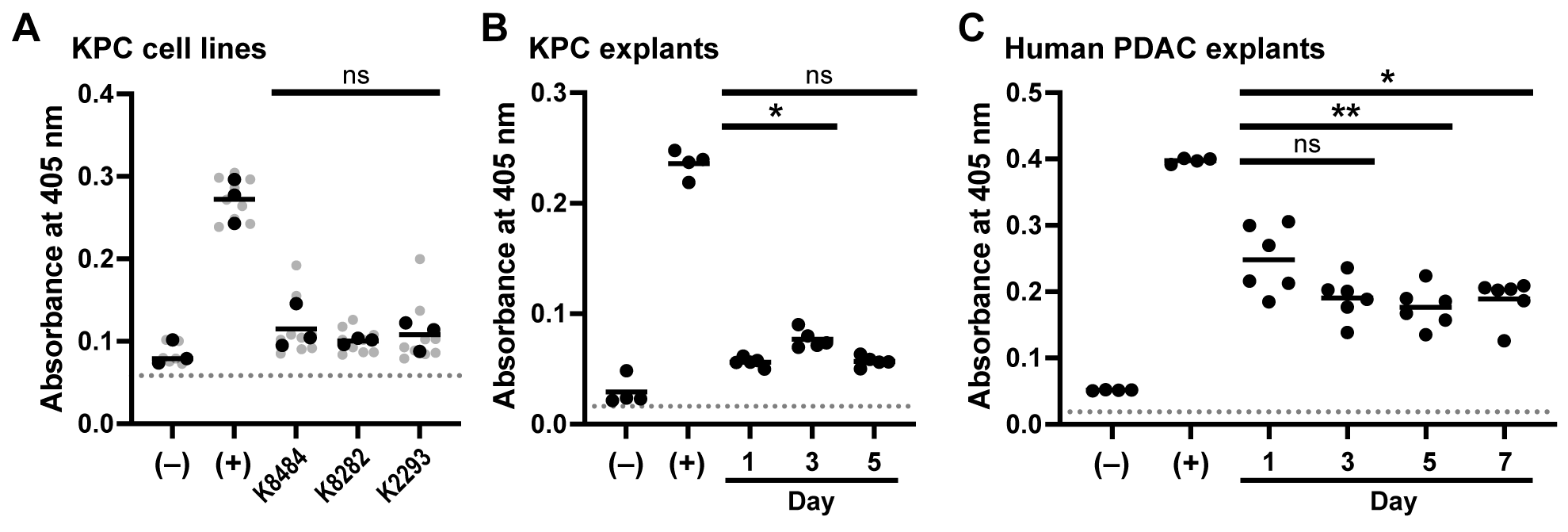
SHH secretion *in vitro* and *ex vivo*. (**A**) SHH secretion was assessed *in vitro* using three KPC-derived tumor cell lines (n=3). As a positive control served SHH-overexpressing Hek293T-SHH cells, negative control is WT Hek293 cells. Dotted line indicates detection threshold. Technical replicates (light shade) are overlaid with average of each biological replicate (dark shade). Statistical analysis was completed using ANOVA with Tukey correction for multiple comparisons. (**B**) SHH secretion in KPC explants over time (n=5). As a positive control served SHH-overexpressing Hek293T-SHH cells, negative control is WT Hek293 cells (n=4). Dotted line indicates detection threshold. Statistical analysis was completed using ANOVA with Tukey correction for multiple comparisons. Significance indicated (*, p<0.05). (**C**) SHH secretion in human PDAC explants over time (n=6). As a positive control served SHH-overexpressing Hek293T-SHH cells, negative control is WT Hek293 cells (n=4). Dotted line indicates detection threshold. Statistical analysis was completed using ANOVA with Tukey correction for multiple comparisons. Significance indicated (*, p<0.05; **, p<0.01).

**Supplementary Figure 5.**
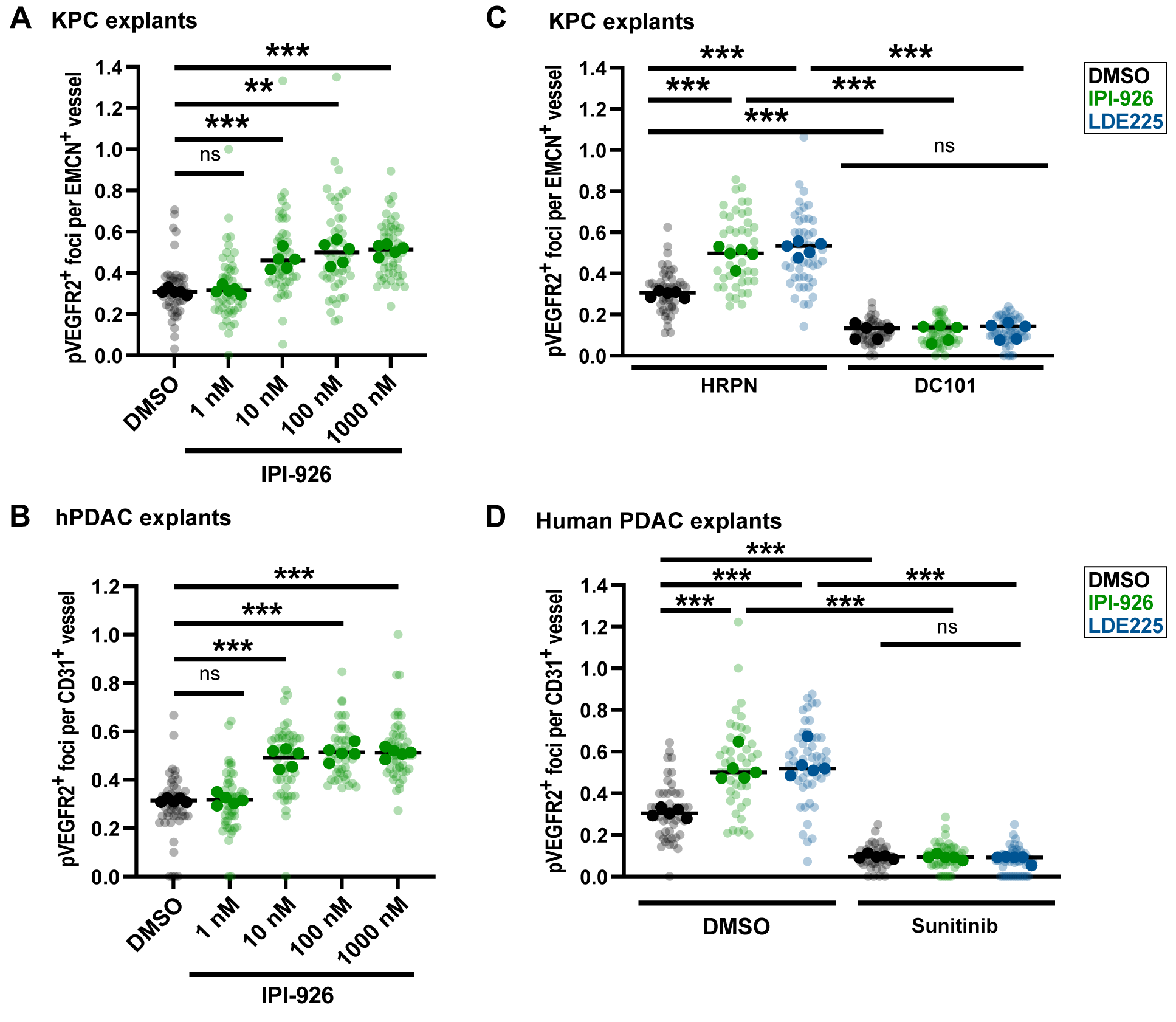
Abrogation of SHH signaling fuels hypersprouting in human and murine PDAC explants. (**A**) Assessment of hypersprouting using co-IF for pVEGFR2/EMCN in KPC explants after 2d of indicated IPI-926 concentrations (n=5). Quantification based on 5-10 fields of view (light shade), averaged per explant (dark shade). Statistical analysis was completed using one-way ANOVA with Tukey correction for multiple comparisons. Significance indicated (**, p<0.01; ***, p<0.001). (**B**) Co-IF for pVEGFR2/CD31 in human PDAC explants after 2d of indicated IPI-926 concentrations (n=5). Quantification based on 5-10 fields of view (light shade), averaged per explant (dark shade). Statistical analysis was completed using one-way ANOVA with one-way Tukey correction for multiple comparisons. Significance indicated (***, p<0.001).(**C**) Co-IF for pVEGFR2/EMCN in KPC explants treated with non-targeting antibodies (HRPN) vs. VEGFR-depleting antibody (DC101) for 2d *ex vivo*. Quantification based on 5-10 fields of view (light shade), averaged per explant (dark shade). Statistical analysis was completed using two-way ANOVA with Dunnett’s correction for multiple comparisons. Significance indicated (***, p<0.001). (**D**) Co-IF for pVEGFR2/EMCN in human PDAC explants treated with DMSO vs. small molecule receptor tyrosine kinase inhibitor sunitinib for 2d *ex vivo*. Quantification based on 5-10 fields of view (light shade), averaged per explant (dark shade). Statistical analysis was completed using two-way ANOVA with Dunnett’s correction for multiple comparisons. Significance indicated (***, p<0.001).

**Supplementary Figure 6.**
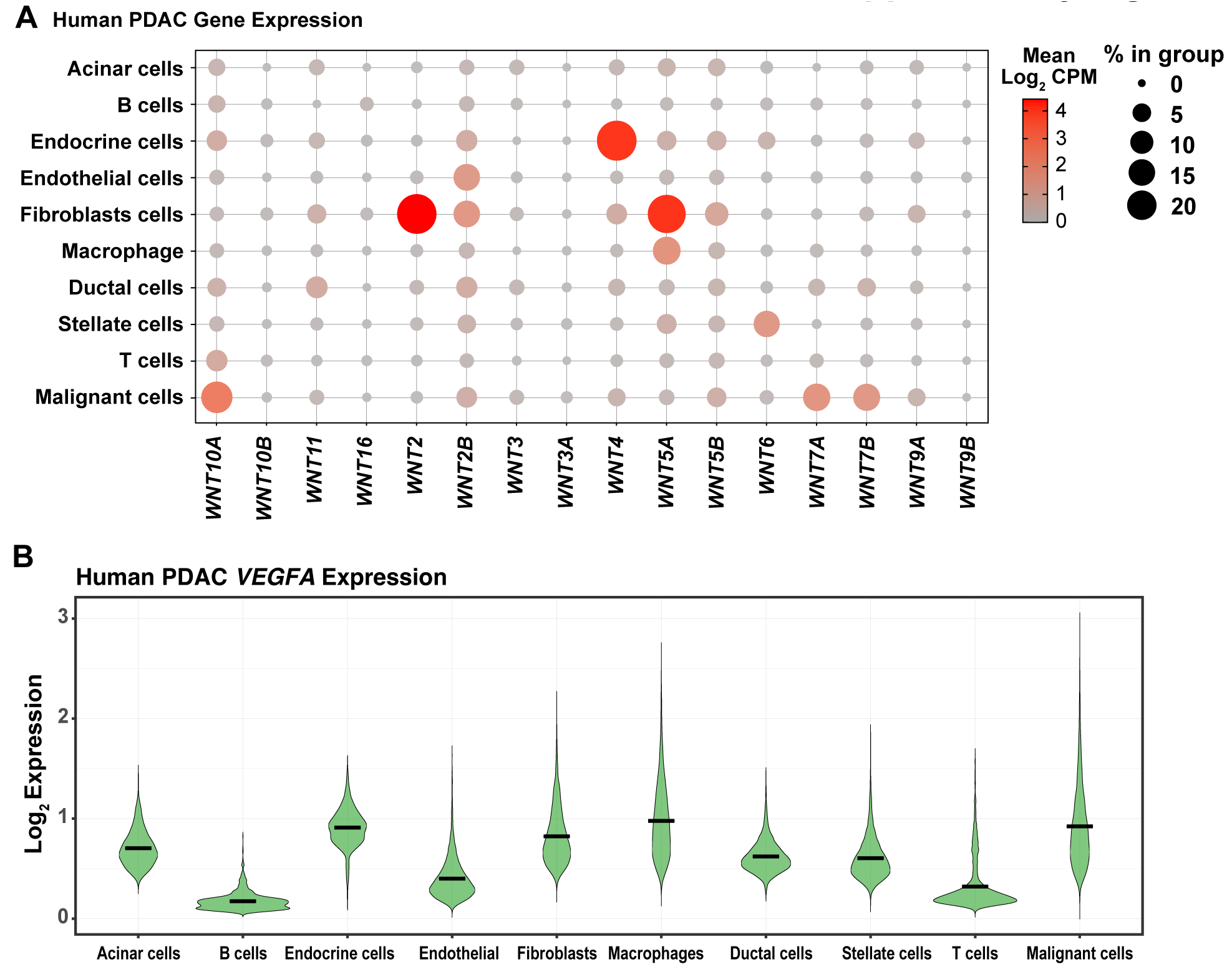
*WNT* and *VEGFA* Expression in human PDAC. (**A**) WNT expression of different cell types in previously published human PDAC scRNA-seq data (48). Only detected WNTs are shown (16 out of 19 known mammalian WNTs). (**B**) *VEGFA* expression of different cell types in previously published human PDAC scRNA-seq data (48).

**Supplementary Figure 7.**
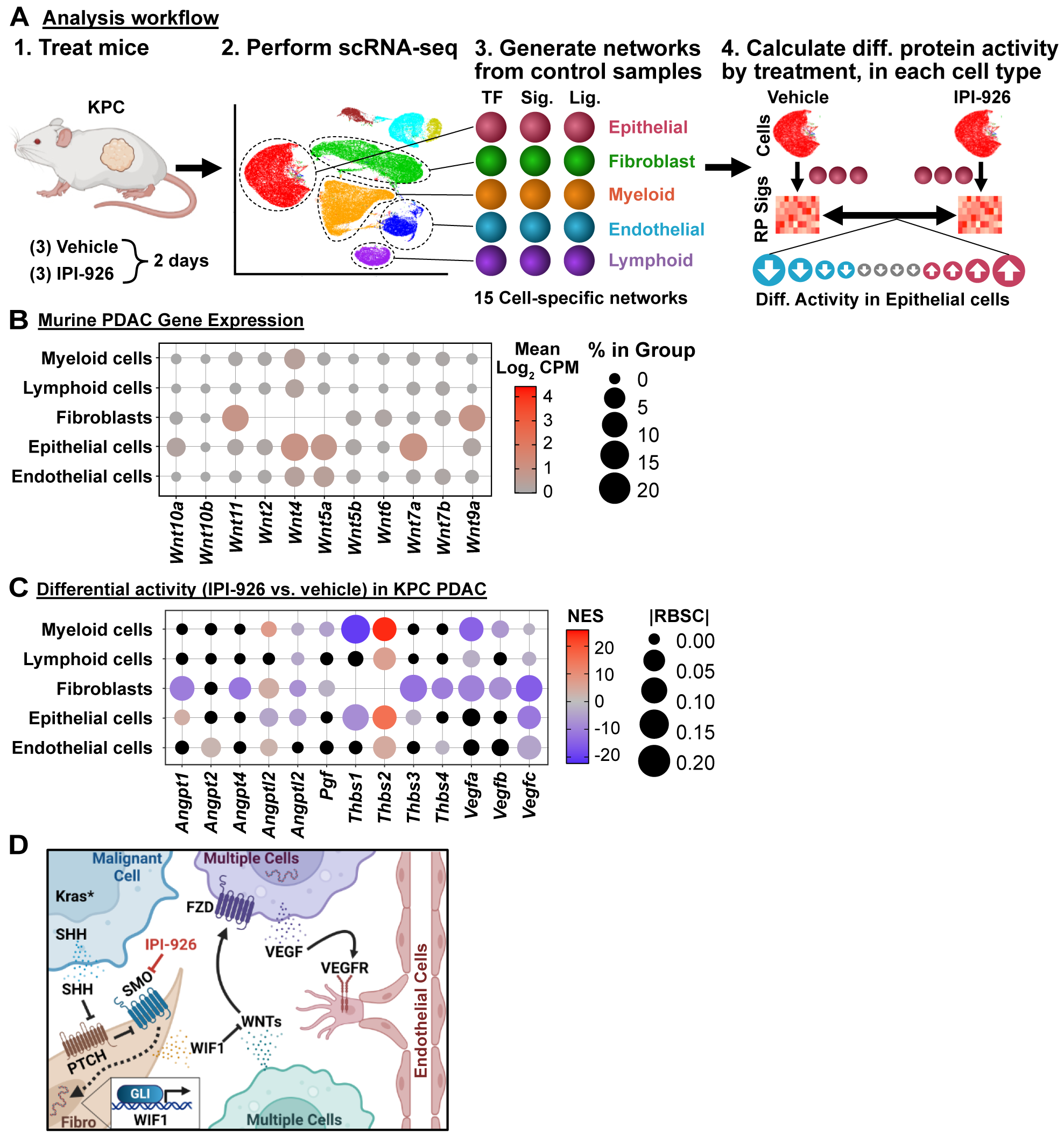
Single cell analysis of paracrine cascades in KPC PDAC. (**A**) Analysis workflow for computational experiments in Fig. 5. **1.** PDAC-bearing KPC mice were treated with vehicle (HPBCD) or IPI-926 for 2 days. **2.** scRNA-seq was performed on each KPC tumor and cell types were clustered by expression. **3.** Three regulatory networks were generated for each major cell type using the cells from vehicle-treated tumors: networks for transcription factors (TFs), signaling proteins (Sig.), and ligands (Lig.). **4.** For each cell types, the three networks were used to calculated the differential protein activities for each TF, signaling protein, and ligand in the networks, comparing vehicle-treated cells to IPI-926-treated cells. (**B**) WNT ligand expression in vehicle-treated KPC mice subjected to scRNA-seq (n=3). (**C**) Differential activity of the indicated angiogenesis ligands upon IPI-926 treatment across all cell types. Non-significant changes are displayed in black. |RBSC| = Rank Biserial Correlation, a measure of effect size. (**D**) Summary schematic of SHH-induced angiosuppression via WIF1, WNT ligands, and VEGFA spanning over multiple cell types in PDAC.

## References

1. Neesse A, Michl P, Frese KK, Feig C, Cook N, Jacobetz MA, et al. Stromal biology and therapy in pancreatic cancer. Gut 2011;60(6):861–8 doi 10.1136/gut.2010.226092.

2. Jacobetz MA, Chan DS, Neesse A, Bapiro TE, Cook N, Frese KK, et al. Hyaluronan impairs vascular function and drug delivery in a mouse model of pancreatic cancer. Gut 2013;62(1):112–20 doi 10.1136/gutjnl-2012-302529.

3. Provenzano PP, Cuevas C, Chang AE, Goel VK, Von Hoff DD, Hingorani SR. Enzymatic targeting of the stroma ablates physical barriers to treatment of pancreatic ductal adenocarcinoma. Cancer Cell 2012;21(3):418–29 doi 10.1016/j.ccr.2012.01.007.

4. Olive KP, Jacobetz MA, Davidson CJ, Gopinathan A, McIntyre D, Honess D, et al. Inhibition of Hedgehog signaling enhances delivery of chemotherapy in a mouse model of pancreatic cancer. Science 2009;324(5933):1457–61.

5. Kahn BM, Lucas A, Alur RG, Wengyn MD, Schwartz GW, Li J, et al. The vascular landscape of human cancer. J Clin Invest 2021;131(2) doi 10.1172/JCI136655.

6. Thayer SP, di Magliano MP, Heiser PW, Nielsen CM, Roberts DJ, Lauwers GY, et al. Hedgehog is an early and late mediator of pancreatic cancer tumorigenesis. Nature 2003;425(6960):851–6 doi 10.1038/nature02009.

7. Rhim AD, Oberstein PE, Thomas DH, Mirek ET, Palermo CF, Sastra SA, et al. Stromal elements act to restrain, rather than support, pancreatic ductal adenocarcinoma. Cancer Cell 2014;25(6):735–47 doi 10.1016/j.ccr.2014.04.021.

8. Steele NG, Biffi G, Kemp SB, Zhang Y, Drouillard D, Syu L, et al. Inhibition of Hedgehog Signaling Alters Fibroblast Composition in Pancreatic Cancer. Clin Cancer Res 2021;27(7):2023–37 doi 10.1158/1078-0432.CCR-20-3715.

9. Happ JT, Arveseth CD, Bruystens J, Bertinetti D, Nelson IB, Olivieri C, et al. A PKA inhibitor motif within SMOOTHENED controls Hedgehog signal transduction. Nat Struct Mol Biol 2022;29(10):990–9 doi 10.1038/s41594-022-00838-z.

10. Basso K, Margolin AA, Stolovitzky G, Klein U, Dalla-Favera R, Califano A. Reverse engineering of regulatory networks in human B cells. Nat Genet 2005;37(4):382–90 doi 10.1038/ng1532.

11. Carro MS, Lim WK, Alvarez MJ, Bollo RJ, Zhao X, Snyder EY, et al. The transcriptional network for mesenchymal transformation of brain tumours. Nature 2010;463(7279):318–25 doi 10.1038/nature08712.

12. Aytes A, Mitrofanova A, Lefebvre C, Alvarez MJ, Castillo-Martin M, Zheng T, et al. Cross-Species Regulatory Network Analysis Identifies a Synergistic Interaction between FOXM1 and CENPF that Drives Prostate Cancer Malignancy. Cancer Cell 2014;25(5):638–51 doi 10.1016/j.ccr.2014.03.017.

13. Ding H, Douglass EF, Jr., Sonabend AM, Mela A, Bose S, Gonzalez C, et al. Quantitative assessment of protein activity in orphan tissues and single cells using the metaVIPER algorithm. Nat Commun 2018;9(1):1471 doi 10.1038/s41467-018-03843-3.

14. Vlahos L, Obradovic A, Worley J, Tan X, Howe A, Laise P, et al. Systematic, Protein Activity-based Characterization of Single Cell State. bioRxiv 2023:2021.05.20.445002 doi 10.1101/2021.05.20.445002.

15. Kerekes K, Banyai L, Patthy L. Wnts grasp the WIF domain of Wnt Inhibitory Factor 1 at two distinct binding sites. FEBS Lett 2015;589(20 Pt B):3044–51 doi 10.1016/j.febslet.2015.08.031.

16. Banyai L, Kerekes K, Patthy L. Characterization of a Wnt-binding site of the WIF-domain of Wnt inhibitory factor-1. FEBS Lett 2012;586(19):3122–6 doi 10.1016/j.febslet.2012.07.072.

17. Hsieh JC, Kodjabachian L, Rebbert ML, Rattner A, Smallwood PM, Samos CH, et al. A new secreted protein that binds to Wnt proteins and inhibits their activities. Nature 1999;398(6726):431–6 doi 10.1038/18899.

18. Mani M, Carrasco DE, Zhang Y, Takada K, Gatt ME, Dutta-Simmons J, et al. BCL9 promotes tumor progression by conferring enhanced proliferative, metastatic, and angiogenic properties to cancer cells. Cancer Res 2009;69(19):7577–86 doi 10.1158/0008-5472.CAN-09-0773.

19. Phng LK, Potente M, Leslie JD, Babbage J, Nyqvist D, Lobov I, et al. Nrarp coordinates endothelial Notch and Wnt signaling to control vessel density in angiogenesis. Dev Cell 2009;16(1):70–82 doi 10.1016/j.devcel.2008.12.009.

20. Kikuchi R, Nakamura K, MacLauchlan S, Ngo DT, Shimizu I, Fuster JJ, et al. An antiangiogenic isoform of VEGF-A contributes to impaired vascularization in peripheral artery disease. Nat Med 2014;20(12):1464–71 doi 10.1038/nm.3703.

21. Huang CL, Liu D, Nakano J, Ishikawa S, Kontani K, Yokomise H, et al. Wnt5a expression is associated with the tumor proliferation and the stromal vascular endothelial growth factor--an expression in non-small-cell lung cancer. J Clin Oncol 2005;23(34):8765–73 doi 10.1200/JCO.2005.02.2871.

22. Peluso MO, Campbell VT, Harari JA, Tibbitts TT, Proctor JL, Whitebread N, et al. Impact of the Smoothened inhibitor, IPI-926, on smoothened ciliary localization and Hedgehog pathway activity. PLoS One 2014;9(3):e90534 doi 10.1371/journal.pone.0090534.

23. Gerhardt H, Golding M, Fruttiger M, Ruhrberg C, Lundkvist A, Abramsson A, et al. VEGF guides angiogenic sprouting utilizing endothelial tip cell filopodia. J Cell Biol 2003;161(6):1163–77 doi 10.1083/jcb.200302047.

24. Sastra SA, Olive KP. Quantification of murine pancreatic tumors by high-resolution ultrasound. Methods Mol Biol 2013;980:249–66 doi 10.1007/978-1-62703-287-2_13.

25. Sastra SA, Olive KP. Acquisition of mouse tumor biopsies through abdominal laparotomy. Cold Spring Harb Protoc 2014;2014(1):47–56 doi 10.1101/pdb.prot077834.

26. Arenas E. Wnt signaling in midbrain dopaminergic neuron development and regenerative medicine for Parkinson’s disease. J Mol Cell Biol 2014;6(1):42–53 doi 10.1093/jmcb/mju001.

27. Bailey JM, Swanson BJ, Hamada T, Eggers JP, Singh PK, Caffery T, et al. Sonic hedgehog promotes desmoplasia in pancreatic cancer. Clin Cancer Res 2008;14(19):5995–6004.

28. Jiang X, Seo YD, Chang JH, Coveler A, Nigjeh EN, Pan S, et al. Long-lived pancreatic ductal adenocarcinoma slice cultures enable precise study of the immune microenvironment. Oncoimmunology 2017;6(7):e1333210 doi 10.1080/2162402X.2017.1333210.

29. Misra S, Moro CF, Del Chiaro M, Pouso S, Sebestyen A, Lohr M, et al. Ex vivo organotypic culture system of precision-cut slices of human pancreatic ductal adenocarcinoma. Sci Rep 2019;9(1):2133 doi 10.1038/s41598-019-38603-w.

30. Kokkinos J, Sharbeen G, Haghighi KS, Ignacio RMC, Kopecky C, Gonzales-Aloy E, et al. Ex vivo culture of intact human patient derived pancreatic tumour tissue. Sci Rep 2021;11(1):1944 doi 10.1038/s41598-021-81299-0.

31. Curiel-Garcia A, Decker-Farrell AR, Sastra SA, Olive KP. Generation of orthotopic patient-derived xenograft models for pancreatic cancer using tumor slices. STAR Protoc 2022;3(4):101899 doi 10.1016/j.xpro.2022.101899.

32. Tremblay MR, Nesler M, Weatherhead R, Castro AC. Recent patents for Hedgehog pathway inhibitors for the treatment of malignancy. Expert Opin Ther Pat 2009;19(8):1039–56 doi 10.1517/13543770903008551.

33. Peng J, Sun BF, Chen CY, Zhou JY, Chen YS, Chen H, et al. Single-cell RNA-seq highlights intra-tumoral heterogeneity and malignant progression in pancreatic ductal adenocarcinoma. Cell Res 2019;29(9):725–38 doi 10.1038/s41422-019-0195-y.

34. Griffin AT, Vlahos LJ, Chiuzan C, Califano A. An Information Theoretic Framework for Protein Activity Measurement. bioRxiv 2021:2021.10.02.462873 doi 10.1101/2021.10.02.462873.

35. Maiti A, Qi Q, Peng X, Yan L, Takabe K, Hait NC. Class I histone deacetylase inhibitor suppresses vasculogenic mimicry by enhancing the expression of tumor suppressor and anti-angiogenesis genes in aggressive human TNBC cells. Int J Oncol 2019;55(1):116–30 doi 10.3892/ijo.2019.4796.

36. Jubb AM, Hurwitz HI, Bai W, Holmgren EB, Tobin P, Guerrero AS, et al. Impact of vascular endothelial growth factor-A expression, thrombospondin-2 expression, and microvessel density on the treatment effect of bevacizumab in metastatic colorectal cancer. J Clin Oncol 2006;24(2):217–27 doi 10.1200/JCO.2005.01.5388.

37. Du W, Menjivar RE, Donahue KL, Kadiyala P, Velez-Delgado A, Brown KL, et al. WNT signaling in the tumor microenvironment promotes immunosuppression in murine pancreatic cancer. J Exp Med 2023;220(1) doi 10.1084/jem.20220503.

38. Sullivan MR, Danai LV, Lewis CA, Chan SH, Gui DY, Kunchok T, et al. Quantification of microenvironmental metabolites in murine cancers reveals determinants of tumor nutrient availability. Elife 2019;8 doi 10.7554/eLife.44235.

39. Ronaldson-Bouchard K, Baldassarri I, Tavakol DN, Graney PL, Samaritano M, Cimetta E, et al. Engineering complexity in human tissue models of cancer. Adv Drug Deliv Rev 2022;184:114181 doi 10.1016/j.addr.2022.114181.

40. Sun H, Li H, Yan J, Wang X, Xu M, Wang M, et al. Loss of CLDN5 in podocytes deregulates WIF1 to activate WNT signaling and contributes to kidney disease. Nat Commun 2022;13(1):1600 doi 10.1038/s41467-022-29277-6.

41. Schindelin J, Arganda-Carreras I, Frise E, Kaynig V, Longair M, Pietzsch T, et al. Fiji: an open-source platform for biological-image analysis. Nat Methods 2012;9(7):676–82 doi 10.1038/nmeth.2019.

42. McCarthy DJ, Chen Y, Smyth GK. Differential expression analysis of multifactor RNA-Seq experiments with respect to biological variation. Nucleic Acids Res 2012;40(10):4288–97 doi 10.1093/nar/gks042.

43. Safgren SL, Olson RLO, Vrabel AM, Almada LL, Marks DL, Hernandez-Alvarado N, et al. The transcription factor GLI1 cooperates with the chromatin remodeler SMARCA2 to regulate chromatin accessibility at distal DNA regulatory elements. J Biol Chem 2020;295(26):8725–35 doi 10.1074/jbc.RA120.013268.

44. Marciniak A, Selck C, Friedrich B, Speier S. Mouse pancreas tissue slice culture facilitates long-term studies of exocrine and endocrine cell physiology in situ. PLoS One 2013;8(11):e78706 doi 10.1371/journal.pone.0078706.

45. Boj SF, Hwang CI, Baker LA, Chio, II, Engle DD, Corbo V, et al. Organoid models of human and mouse ductal pancreatic cancer. Cell 2015;160(1-2):324–38 doi 10.1016/j.cell.2014.12.021.

46. Stuart T, Butler A, Hoffman P, Hafemeister C, Papalexi E, Mauck WM, 3rd, et al. Comprehensive Integration of Single-Cell Data. Cell 2019;177(7):1888–902 e21 doi 10.1016/j.cell.2019.05.031.

47. Hafemeister C, Satija R. Normalization and variance stabilization of single-cell RNA-seq data using regularized negative binomial regression. Genome Biol 2019;20(1):296 doi 10.1186/s13059-019-1874-1.

48. Tickle T, Tirosh I, Georgescu C, Brown M, Haas B. 2019 inferCNV of the Trinity CTAT Project. <https://github.com/broadinstitute/inferCNV>.

49. Mann HB, Whitney DR. On a Test of Whether one of Two Random Variables is Stochastically Larger than the Other. The Annals of Mathematical Statistics 1947;18(1):50–60, 11.

50. Benjamini Y, Hochberg Y. Controlling the False Discovery Rate: A Practical and Powerful Approach to Multiple Testing. Journal of the Royal Statistical Society Series B (Methodological) 1995;57(1):289–300.

51. Gu Z, Eils R, Schlesner M. Complex heatmaps reveal patterns and correlations in multidimensional genomic data. Bioinformatics 2016;32(18):2847–9 doi 10.1093/bioinformatics/btw313.

52. Wickham H. ggplot2: Elegant Graphics for Data Analysis.: Springer-Verlag New York; 2016.

